# Heterogeneous and Novel Transcript Expression in Single Cells of Patient-Derived ccRCC Organoids

**DOI:** 10.1101/2024.03.15.585271

**Authors:** Tülay Karakulak, Natalia Zajac, Hella Anna Bolck, Anna Bratus-Neuenschwander, Qin Zhang, Weihong Qi, Debleena Basu, Tamara Carrasco Oltra, Hubert Rehrauer, Christian von Mering, Holger Moch, Abdullah Kahraman

**Author notes:** shared first author (Tülay Karakulak, Natalia Zajac).

## Abstract

Splicing is often dysregulated in cancer, leading to alterations in the expression of canonical and alternative splice isoforms. This complex phenomenon can be revealed by an in-depth understanding of cellular heterogeneity at the single-cell level. Recent advances in single-cell long-read sequencing technologies enable comprehensive transcriptome sequencing at the single-cell level. In this study, we have generated single-cell long-read sequencing of Patient-Derived Organoid (PDO) cells of clear-cell Renal Cell Carcinoma (ccRCC), an aggressive and lethal form of cancer that arises in kidney tubules. We have used the Multiplexed Arrays Sequencing (MAS-ISO-Seq) protocol of PacBio to sequence full-length transcripts exceptionally deep across 2,599 single cells to obtain the most comprehensive view of the alternative landscape of ccRCC to date. On average, we uncovered 86,182 transcripts across PDOs, of which 31,531 (36.6%) were previously uncharacterized. In contrast to known transcripts, many of these novel isoforms appear to exhibit cell-specific expression. Nonetheless, >50% of these novel transcripts were predicted to possess a complete protein-coding open reading frame. This finding suggests a biological role for these transcripts within kidney cells. Moreover, an analysis of the most dominant transcript switching events between ccRCC and non-ccRCC cells revealed that many switching events were cell and sample-specific, underscoring the heterogeneity of alternative splicing events in ccRCC.

Overall, our research elucidates the intricate transcriptomic architecture of ccRCC, potentially exposing the mechanisms underlying its aggressive phenotype and resistance to conventional cancer therapies.

## Introduction

Alternative splicing is a pivotal mechanism by which eukaryotic cells enhance their transcriptomic and proteomic diversity (Graveley 2001). By allowing a single gene to encode multiple RNA variants, alternative splicing contributes significantly to cellular complexity, tissue specificity (Xu 2002), and organismal adaptability (Marasco and Kornblihtt 2023; Verta and Jacobs 2022). In the context of human disease, notably cancer, dysregulation of alternative splicing events can lead to the expression of oncogenic isoforms, influencing tumor initiation, progression, and resistance to therapy (Sciarrillo et al. 2020; Bradley and Anczuków 2023). Despite its recognized importance, the comprehensive characterization of alternative splicing at the resolution of individual cells remains a formidable challenge, primarily due to the limitations of conventional sequencing technologies in capturing the full spectrum of splicing events.

Recent advances in single-cell RNA sequencing (scRNA-seq) have revolutionized our understanding of cellular heterogeneity in complex tissues and tumoral environments, revealing unprecedented insights into the transcriptomic variations that define cell types, states, and functions (Cha and Lee 2020; Li et al. 2022; Travaglini et al. 2020; Yao et al. 2023; Dondi et al. 2023). However, most single-cell studies have relied on short-read sequencing technologies, which, despite their high throughput, fall short of accurately resolving complex splice variants due to their limited read lengths. Long-read sequencing technologies offer a promising solution to these limitations. With the ability to generate reads that span entire transcript isoforms, long-read sequencing enables the direct observation of splicing patterns and the identification of novel isoforms that would be missed or misassembled by short-read technologies (Byrne et al. 2017; Amarasinghe et al. 2020; Bolisetty et al. 2015). However, long-read sequencing was not appropriate for single-cell transcriptome measurements due to the initial lower throughput and high sequencing errors. With the recent advances in sequencing chemistries and transcript concatenation protocols, the restrictions could be overcome, allowing us to measure transcripts in the transcriptome at full-length at single-cell resolution.

Using derivatives of this new technology, the research community has begun to investigate the transcriptome of various samples at single-cell resolution. For example, Shiau *et al*. identified a distinct combination of isoforms in tumor and neighboring stroma/immune cells in a kidney tumor, as well as cell-type-specific mutations like VEGFA mutations in tumor cells and HLA-A mutations in immune cells (Shiau et al. 2023). Tian *et al*. highlighted the complexity of the transcriptome in human and mouse samples by identifying thousands of novel transcripts with conserved functional modules enriched in alternative transcript usage, including ribosome biogenesis and mRNA splicing. They found drug-resistance mutations in subclones within transcriptional clusters (Tian et al. 2021). Also, Yang *et al*. observed thousands of novel transcripts in human cerebral organoids, with differentially spliced exons and retained introns (Yang et al. 2023). Cell-type-specific exons with de novo mutations were enriched in autistic patients. In another interesting study, Wan *et al*. integrated single-cell long-read sequencing with single-molecule microscopy and observed distinct but consistent bursting expression for all genes with similar nascent RNA dwell time (Wan et al. 2021; Shiau et al. 2023). The intron removal time spans minutes to hours, suggesting that the spliceosome removes introns progressively in pieces. In a recent study, Dondi *et al*. identified over 52,000 novel transcripts in five ovarian cancer samples that had not been reported previously, and similar to the studies above, discovered cell-specific transcript and polyadenylation site usages and were able to identify a gene fusion event that would have been missed using short-read sequencing (Dondi et al. 2023).

Clear cell renal cell carcinoma (ccRCC) is the most prevalent form of kidney cancer, characterized by its heterogeneous cellular composition and a complex genetic landscape (Hsieh et al. 2017; Turajlic et al. 2018a). A hallmark of ccRCC is the loss of the von Hippel-Lindau (VHL) tumor suppressor gene through genetic (point mutations, indels and 3p25 loss) and/or epigenetic (promoter methylation) mechanisms. Loss of the VHL gene can lead to the stabilization of hypoxia- inducible factors (HIFs) and subsequent activation of an hypoxic response even in oxygenated tissue microenvironment. The resulting uncontrolled activation of transcriptional targets that regulate angiogenesis, metabolic pathways, apoptosis, and other processes can drive tumor progression and survival while inducing the acceleration of clonal evolution and subclonal diversification (Turajlic et al., 2018).

Here, we have applied PacBio’s new Multiplexed Arrays Sequencing (MAS-ISO-Seq) protocol (Al’Khafaji et al. 2023) to probe full-length transcriptomic profiles of single cells in patient-derived kidney organoids from four individuals with ccRCC. Importantly, the MAS-ISO-Seq method hinges on the availability of intact RNA molecules that can exclusively be obtained from viable cells, preventing its application for archival formalin-fixed paraffin embedded samples. Cancer-derived organoids serve as an ideal starting material. They closely mirror important biological features of the original tumors including genetic intra-tumor heterogeneity (ITH) as a three- dimensional model, and provide a renewable source of living cells for analysis (Bolck et al. 2021). Thus, our organoids allow for an unprecedented exploration of the transcriptomic diversity within patient-derived cellsand reveal important insights into the mechanisms driving tumor evolution and therapy resistance. By applying long-read single-cell RNA sequencing to PDOs, one can derive important insights into the transcriptional landscapes of these important translational models.

Despite extensive research highlighting the role of alternative splicing in ccRCC development and treatment response (Wang et al. 2022; Simmler et al. 2022; Zhang et al. 2021a), the transcriptome landscape of ccRCC at the single-cell resolution remains unexplored. Given the well-known genetic heterogeneity and complexity of the tumor microenvironment in ccRCC (Turajlic et al. 2018a, 2018b), understanding these processes at the single-cell resolution could reveal critical insight for ccRCC biology. For example, recent single-cell studies have suggested VCAM1- positive renal proximal tubule cells to be the likely origin of ccRCC (Zhang et al. 2021b; Schreibing and Kramann 2022), which is consistent with the hypothesis that ccRCC is derived from the proximal tubules. Also, ccRCC tumors were found to detain many CD8+ T-cells and macrophages in immune checkpoint inhibition responsive and resistant samples, respectively (Krishna et al. 2021). The distinct response could explain the general good response of ccRCC patients to immunotherapy despite having a low mutational burden in their ccRCC tumors (Borcherding et al. 2021).

Here, for the first time, we explored the transcriptome landscape of ccRCC samples and one matched-normal patient-derived organoids (PDOs) in single-cell resolution using single-cell long-read sequencing technology. We aimed to understand the heterogeneity within cells and between samples at the alternative splicing level, and identify isoform switching events in ccRCC cells that could pave the way for novel therapeutic strategies.

## Results

### Full-length single-cell sequencing reveals transcript diversity and the cell heterogeneity of known and novel transcripts

To discern the transcriptome diversity in ccRCC, we have applied full-length single-cell sequencing using the MAS-Seq protocol (Al’Khafaji et al. 2023) on a PacBio Sequel IIe instrument to five patient- derived organoids (PDO) samples (Fig. 1A).The PDOs were established from fresh tissue samples obtained from four individuals with ccRCC (Fig. 1B). We included one PDO that was generated from matching normal kidney tissue from sample ccRCC2. All ccRCC-derived organoids carried a VHL mutation, a hallmark of ccRCC (Table 1). To sequence the single-cell transcriptomes as deeply as possible, we loaded transcript molecules of as few cells as possible on the flow cell. With 29.4 to 58.8 million segmented reads per sample we sequenced 310 - 1091 cells and obtained a total of 216,926 - 346,107 transcripts. The average sequencing depth thus ranged from 21,499 to 96,620 reads per cell (Table 2). Calculation of the number of unique genes and transcripts and their UMI counts per cell revealed that the ccRCC4 PDO with the highest number of cells had the lowest number of transcripts, genes, and UMI per cell (Table 2, Figure 1C, Supplementary Fig. 1A).

**Fig 1.**
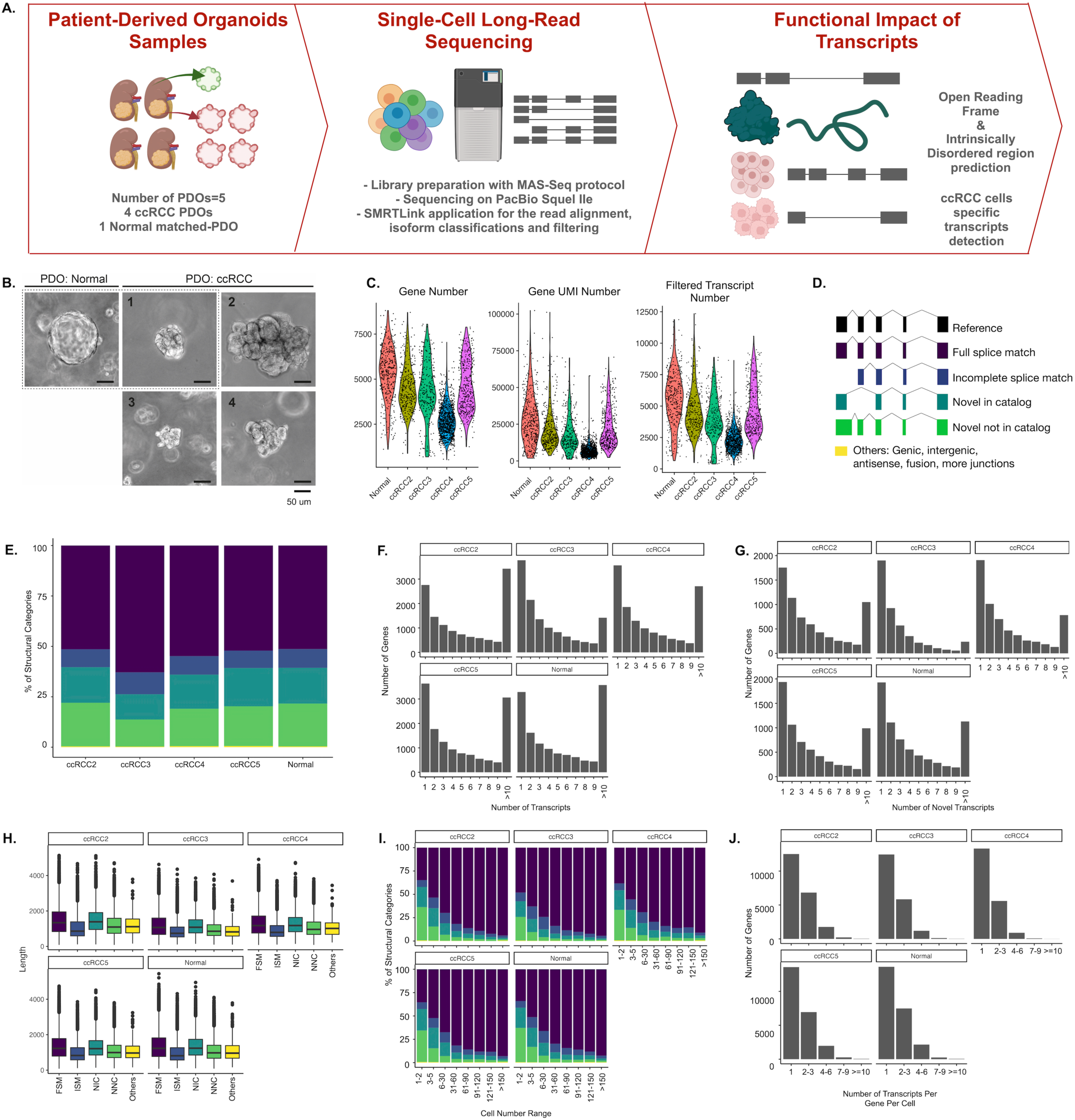
Transcript landscape and cell heterogeneity in Normal and ccRCC-PDOs: **(A)** Schematic design of the project showing how patient-derived organoid (PDO) samples are established, sequenced using single-cell long-read sequencing, and functionally characterized (illustrations were created by Biorender (BioRender.com)). **(B)** The brightfield representative images of our organoids. The dotted line marks the matched pair. The scale bar is 50um. **(C).** Distribution of number of genes, UMI counts and filtered transcript numbers across samples. (D). SQANTI3 transcript categories. **(E).** The proportion of transcript categories found across four ccRCC-PDO and one normal PDO (see the (C) for the color code). **(F).** The number of identified transcripts per gene in each PDO. The x-axis denotes the number of transcripts per gene, categorized into bins (1, 2, 3, 4, 5, 6, 7, 8, 9, and >10), while the y-axis represents the number of genes. The height of each bar reflects the count of genes that express the corresponding number of transcripts. **(G).** The number of identified novel transcripts per gene per cell in each PDO. The x-axis shows the number of transcripts detected per gene per cell, categorized into different bins, while the y-axis denotes the total number of genes, with the height of each bar reflecting the count of genes that express the corresponding number of transcripts per cell. **(H).** Distribution of transcript lengths for each structural category across samples. **(I).** Proportional distribution of identified transcripts’ structural categories across cell number ranges. **(J).** Number of transcripts per gene per cell across samples, categorized into bins (1, 2-3, 4-6, 7-9, and >10).

**Table 1:**
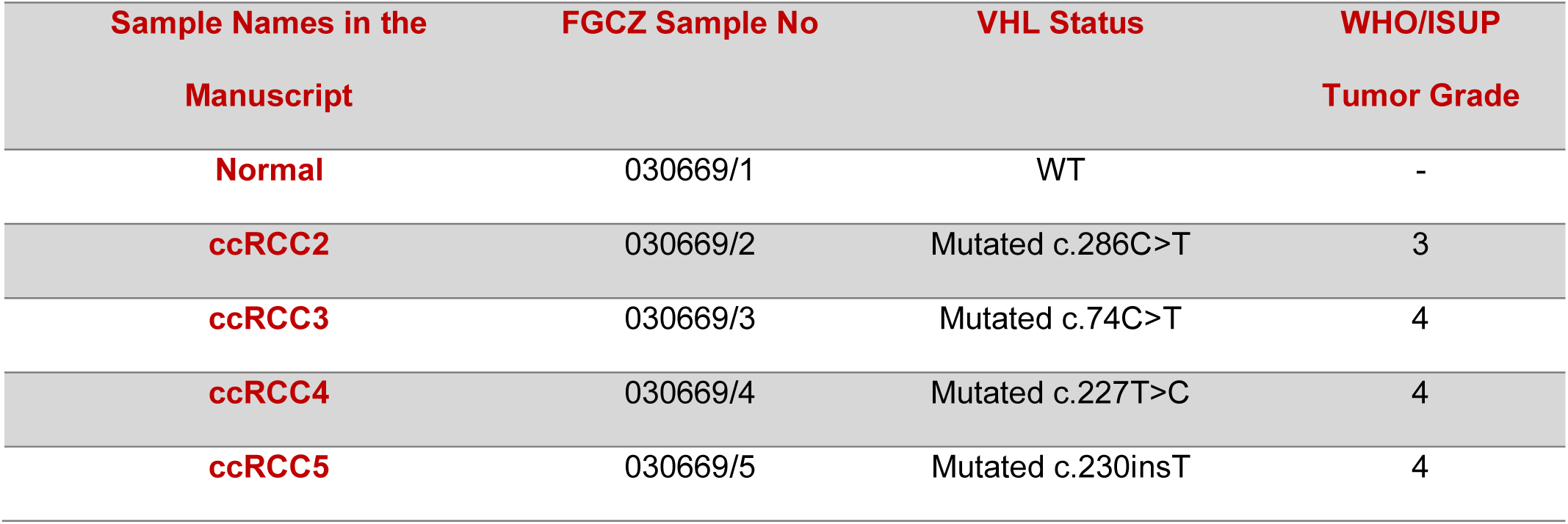
Clinical data of patient-derived organoid (PDO) samples.

| Sample Names in the<br>Manuscript | FGCZ Sample No | VHL Status | WHO/ISUP<br>Tumor Grade |
| --- | --- | --- | --- |
| Normal | 030669/1 | WT | - |
| ccRCC2 | 030669/2 | Mutated c.286C>T | 3 |
| ccRCC3 | 030669/3 | Mutated c.74C>T | 4 |
| ccRCC4 | 030669/4 | Mutated c.227T>C | 4 |
| ccRCC5 | 030669/5 | Mutated c.230insT | 4 |

**Table 2:** The number of HiFi reads, segmented reads, cells, genes, and transcripts and their structural categories before and after filtering. **Number of genes, transcripts, and the structural categories after filtering. FSM: Full splice match, ISM: Incomplete splice match, NIC: Novel In Catalog, NNC: Novel Not In Catalog. Other: Genic, antisense, intergenic, fusion, more Junctions.

|  | Normal | ccRCC2 | ccRCC3 | ccRCC4 | ccRCC5 |
| --- | --- | --- | --- | --- | --- |
| Cells | 437 | 373 | 310 | 1091 | 388 |
| HiFi Reads | 3,504,085 | 3,785,895 | 2,166,307 | 1,880,180 | 2,275,670 |
| Segmented reads (S-reads) | 54,962,298 | 58,333,415 | 34,180,213 | 29,404,003 | 35,635,073 |
| Mean length of S-reads | 844.2 | 910.0 | 801.0 | 859.0 | 890.0 |
| Reads after Barcode Correction and UMI Deduplication | 27,799,824 | 18,554,804 | 15,850,578 | 21,075,848 | 19,102,522 |
| Reads in cells | 68% | 65% | 68% | 83% | 65% |
| Mean reads per cell | 82,703 | 96,620 | 70,423 | 21,499 | 57,006 |
| Median UMIs per cell | 56437.0 | 41881.5 | 42798.5 | 17705.0 | 40062.5 |
| Unique Genes | 29,138 | 26,260 | 26,657 | 29,074 | 30,065 |
| Unique Transcripts | 346,107 | 303,547 | 216,926 | 289,556 | 301,160 |
| FSM (%) | 71,205<br>(20.5%) | 68,352<br>(22.5%) | 50,983<br>(23.5%) | 62,582<br>(21.6%) | 64,378<br>(21.4%) |
| ISM (%) | 126,715<br>(36.6%) | 101,665<br>(33.5%) | 96,380<br>(44.4%) | 109,263<br>(37.7%) | 102,536<br>(34%) |
| NIC (%) | 52,740<br>(15.2%) | 49,142<br>(16.2%) | 23,054<br>(10.6%) | 42,698<br>(14.7%) | 49,049<br>(16.3%) |
| NNC (%) | 83,308<br>(24.1%) | 73,820<br>(24.3%) | 37,997<br>(17.5%) | 62,710<br>(21.7%) | 72,372<br>(24.0%) |
| Other (%) | 12,139<br>(3.5%) | 10,568<br>(3.5%) | 8,512<br>(3.9%) | 12,303<br>(4.2%) | 12,825 (4.3%) |
| Cells** | 390 | 334 | 272 | 1016 | 366 |
| Unique Genes** | 13,630 | 12,532 | 12,460 | 13,323 | 13,586 |
| Unique Transcripts** | 101,942 | 97,294 | 56,596 | 83,342 | 91,734 |
| FSM** (%) | 52,327<br>(51.3%) | 50,011<br>(51.4%) | 35,585<br>(62.9%) | 45,697<br>(54.8%) | 47,774<br>(52.1%) |
| **ISM\*\* (%)** | 9,490<br>(9.31%) | 8,689<br>(8.9%) | 6,154<br>(10.9%) | 7,649<br>(9.2%) | 7,930<br>(8.64%) |
| **NIC\*\* (%)** | 18,084<br>(17.7%) | 17,168<br>(17.6 %) | 7,094<br>(12.5%) | 14,123<br>(16.9%) | 17,406<br>(19.0%) |
| **NNC\*\* (%)** | 21,632<br>(21.2%) | 21,020<br>(21.6%) | 7,591<br>(13.4%) | 15,444<br>(18.5%) | 18,092<br>(19.7%) |
| **Other\*\* (%)** | 409 (0.4%) | 406 (0.417) | 172 (0.3%) | 429 (0.5%) | 532 (0.6%) |

The Iso-seq pipeline classified transcripts into four categories using SQANTI3 in SMRT-Link. Based on the alignment profile of exon coordinates of transcripts to the reference transcriptome, SQANTI3 (Pardo-Palacios et al. 2023) categorized the transcripts as full-splice match (FSM), incomplete- splice match (ISM), novel in catalog (NIC), and novel not in catalog (NNC) (Fig. 1D). FSM transcripts perfectly align with reference transcripts at their junctions; ISM transcripts have fewer exons at the 5’ or 3’ ends, while the rest of the internal junctions align with the reference transcript junctions. The novel transcript categories NIC or NNC are made of new combinations of known splice junctions or have at least one new donor or acceptor site, respectively. In addition, SQANTI3 sub-categorizes isoforms based on their 5’ and 3’ ends (Pardo-Palacios et al. 2023). We grouped the remaining SQANTI3 transcripts, namely antisense, genic intron, genic genomic, and intergenic, into a single category called ’Other’. Filtering based on the CAGE peak, 3’ and 5’ support and TSS ratio left on average 86,182 isoforms, of which 31,531 (36.6%) were novel (Table 2). While 37.2% of the transcripts were identified as ISM before filtering, this number reduced to 9.31% showing that many ISM isoforms have missing 3’ or 5’ support and are prone to degradation. Filtered isoforms have on average 53.7% FSM followed by 36.6% % novel transcripts, of which 17.1% and 19.4% were identified as NIC and NNC, respectively. (Fig. 1E, Table 2).

FSM transcripts mainly consisted of a subcategory of transcripts having alternative 3’ ends, while ISM transcripts of those with alternative 5’ prime ends (Supplementary Figure 1B). We detected more than 10 transcripts for the 26% of genes. ccRCC3 had the smallest number of genes (11%) expressing more than ten transcripts. (Fig. 1F). With 8%, the highest percentage of genes with at least 10 or more novel isoforms were detected in ccRCC2 and Normal samples. (Fig. 1G). FSM and NIC transcripts tended to be the longest and to have similar lengths on average with 1,338 bp length (t-test p-value=0.53) (Fig. 1H), while the ISM transcripts showed shorter lengths compared to FSM and NIC (t-test, p-value < 2.2e-16). On average, ∼50% of transcripts found in only one cell were novel, while 93% of the transcripts found in more than 150 cells were FSM (Fig. 1I). Most genes were found to express one transcript per cell across all samples (Fig. 1J).

Both FSM and NIC transcripts tended to have a higher expression within a cell if they were also expressed in many cells (Supplementary Fig. 1C and 1D). The highest UMI counts were found for FSM transcripts compared to other categories (Supplementary Fig. 1E). Of the novel isoforms, 63.3% were found only within one cell of a sample. However, there were some exceptions, for example, the novel transcripts of the genes Glyceraldehyde 3-phosphate dehydrogenase (*GAPDH)*, Pyruvate kinase (*PKM)*, Angiopoietin-like 4 (*ANGPTL4)*, Nicotinamide N-methyltransferase (NNMT), and Karyopherin Subunit Alpha 2 (KPNA2) were expressed in at least 30% of the cells of one or more samples (see Supplementary File 1 for the full list). Among those, *GAPDH*, *PKM* are known to be involved in glycolysis, and it is reported that enzymes having a role in glycolysis are upregulated in the occurrence of VHL-deficient ccRCC due to the upregulation in hypoxia-inducible factor 1alpha (HIF-1a) (Miranda-Poma et al. 2023). *ANGPTL4* is another hypoxia-inducible gene, and its expression has been shown as a potential diagnostic marker for ccRCC (Verine et al. 2010). KPNA2 is overexpressed in many cancers (Sun et al. 2021) including ccRCC, and its knockdown has been shown to inhibit kidney tumor proliferation (Zheng et al. 2021). *NNMT* is another gene overexpressed in ccRCC, and it was previously characterized as a promising drug target for ccRCC (Reustle et al. 2022). Our findings suggest that those novel transcripts expressed more broadly across cells might play an important role in the pathogenesis of ccRCC.

### More than 50% of novel transcripts have translation capability

To assess whether the novel transcripts are protein coding, we predicted the Open Reading Frame (ORF) using TransDecoder (Haas BJ.). Based on the occurrence of start and stop codons and coding regions, TransDecoder assigned transcripts into varying sub-ORF categories, including 3’ partial (transcripts with missing stop codons), 5’ partial (transcripts with missing start codons), internal (transcripts that miss both start and stop codons), and complete transcripts (including all necessary parts to code a protein) (Figure 1B). For about 77.89% of the NIC transcripts, we were able to predict an ORF (Fig. 2A). Even after applying our stringent filtering criteria, ISM transcripts remained with the lowest proportion of complete ORFs (Fig. 2C). We also investigated the prevalence of sub-ORF categories of novel transcripts across varying cell number ranges. Transcripts commonly expressed in a sample were significantly more likely to have complete ORFs as compared to cell-specific transcripts, (Cochran-Armitage test for trend test: p<2.2e-16) (Fig. 2D). To understand whether the predicted protein isoforms form a stable protein structure that could hint towards a biological function, we predicted intrinsically disordered regions for all isoforms with complete ORFs using iupred2 (Mészáros et al. 2018). The calculations demonstrated that ISM transcripts had the highest proportion of disordered residues (Wilcoxon rank sum test, p.adj: ISM- FSM: 8e-13, ISM-NIC: 2.20e-18, ISM-NNC p-adj value: 4e-21,) (Fig. 2E) and NNC and NIC transcripts with intron retention showed a higher disordered score than those with a new splice site (Wilcox rank sum test p.adj value: <2e-16), combination of known junctions (Wilcoxon rank sum test, p-adj value: 1.7e-14) and known splice sites (Wilcoxon rank sum test, p-adj value: < 2e-16). On the other hand, transcripts with a new combination of a splice site from the NIC category showed the least proportion of disordered regions (Fig. 2F). For example, we identified eight novel transcripts of Nicotinamide-N-methyltransferase (*NNMT*) in ccRCC2 PDO, each comprising three to four exons, and each with a complete ORF. The protein sequences encoded by these transcripts were characterized by more than 88% of their residues being ordered. These transcripts were found to be expressed in a range of 1 to 144 cells. On the other hand, protein sequences of novel ADP Ribosylation Factor Like GTPase 6 Interacting Protein 4 (ARL6IP4) transcripts with complete ORF exhibited on average 94.1% of their residues as disordered.

**Fig 2:**
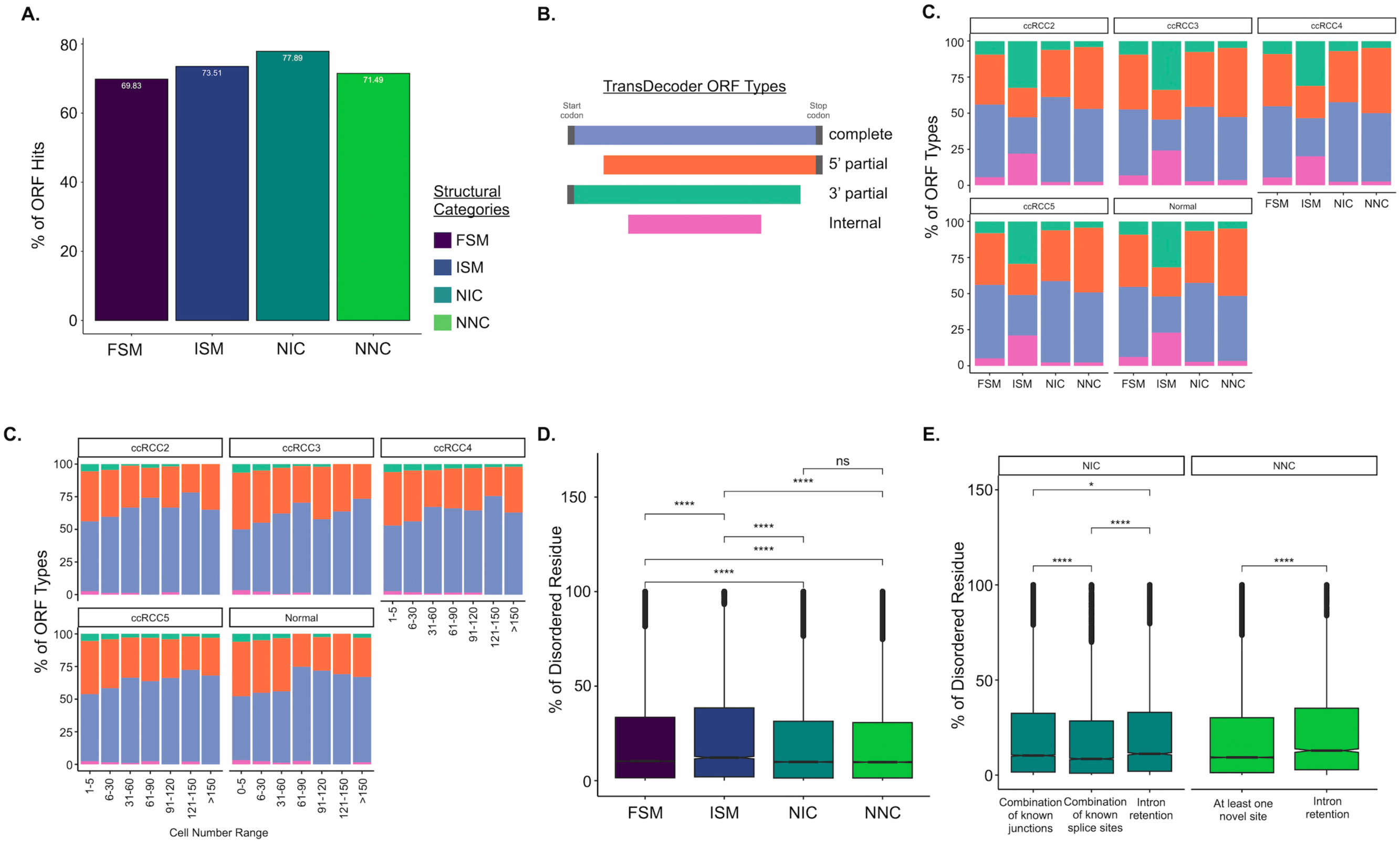
Distribution of open reading frame (ORF) categories and intrinsically disordered protein predictions: (A). Percentage of ORF hits across different structural categories in all datasets. For percentage of ORF hits across isoform subcategories see Supplementary Figure 2A. **(B).** ORF types predicted by Transdecoder. **(C).** The fraction of different ORF types across datasets in each structural category. Each color represents various ORF types. **(D).** Distribution of ORF types in novel transcripts as a function of cell number range across datasets. The x-axis categorizes the cell number range, while the y-axis shows the proportion of each ORF type (see (B) for the legends). **(E).** Comparison of disordered scores for the protein sequences of complete-ORF transcripts across structural categories. **(F).** Comparison of disordered scores for the protein sequence of NIC and NNC showing complete ORFs across different sub-structural categories. For the percentage of disordered region scores across FSM subcategories, see Supplementary Figure 2B.

### Genes expressing ccRCC Cell-Specific Transcripts play a role in ccRCC related pathways

To evaluate the cell types in our samples, we examined the expression of ccRCC-specific marker (CA9), a kidney proximal tubule marker (GGT1), and epithelial cell marker (EPCAM). ccRCC marker CA9 was predominantly expressed in PDO cells from samples ccRCC2, ccRCC4, and ccRCC5 (Fig. 3A, Supplementary Figure 3B, 3D, 3E). As ccRCC originates from the proximal tubule (PTC), we also found that nearly all CA9 expressing cells also expressed the PTC marker GGT1 (Supplementary Figure 3) Interestingly, EPCAM was expressed predominantly in the normal sample and in the PDO cells of ccRCC3 (Supplementary Figure 3A, 3C). In addition, we annotated the cells with a manually curated list of genes and observed that majority of CA9 expressing cells were annotated as ccRCC or Epithelial-Mesenchymal Transition (EMT) process across ccRCC2, ccRCC4, and ccRCC5 (Supplementary Figure 4). The transcript expression profile of ccRCC3 stood out compared to the other ccRCC organoids as we could not detect CA9 expression in this organoid sample. Interestingly, ccRCC3 had P25L at the *VHL* mutations (Table 1). This variant was previously described as a polymorphic likely benign mutation, which could explain the VHL-positive-like expression profile of ccRCC3 (Rothberg 2001; Nickerson et al. 2008) lacking overexpression of the VHL-HIF pathway.

**Fig 3:**
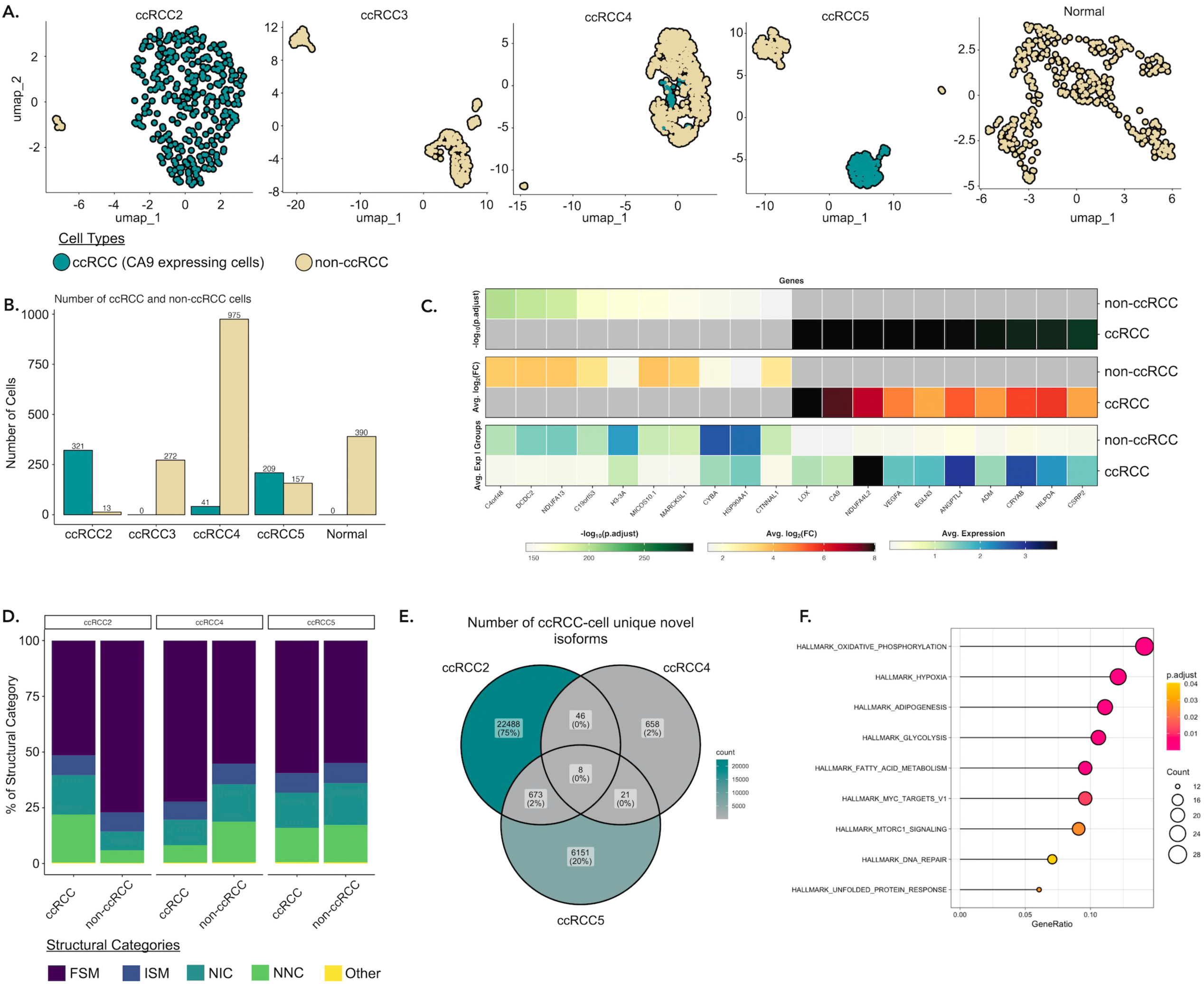
Categorizing cells as ccRCC and non-ccRCC in PDOs: **(A)** UMAP plot of the ccRCC marker CA9 expression in cells across all PDOs, with darker colors indicating CA9 expression levels. **(B).** The table shows the number of ccRCC and non-ccRCC cells in each PDO categorized based on their CA9 expression. **(C).** The heatmap shows the differential gene expression between ccRCC and non-ccRCC cells. **(D).** The proportion of expressed transcripts’ structural categories across ccRCC and non-ccRCC cells. **(E).** Overlap of novel isoforms unique to ccRCC cells only. **(F).** Over-representation hallmark analysis of genes expressing common novel transcripts explicitly in ccRCC cells.

Moreover, to explore the gene and transcript diversity between typical ccRCC and non-ccRCC cells, we categorized cells based on their CA9 expression. CA9 expression is a result of HIF up-regulation due to VHL inactivation (Tostain et al. 2010). ccRCC2, ccRCC4, and ccRCC5 samples contained 321 (96.1%), 41 (4%), and 209 (57%) ccRCC cells with CA9 expression, respectively (Figure 3B). Differential gene expression analysis between ccRCC and non-ccRCC cells revealed upregulation of several ccRCC-related genes in ccRCC cells, including NADH dehydrogenase 1 alpha subcomplex, 4-like 2 (*NDUFA4L2*), Lysyl oxidase (*LOX*), Vascular Endothelial Growth Factor A (*VEGFA*), *ANGPTL4*, and Egl-9 Family Hypoxia Inducible Factor 3 (*EGLN3*) (Fig. 3C). Each of these genes is known to have a role in the progression of ccRCC through various mechanisms. NDUFA4L2 and EGLN3 are critical for the adaptation of ccRCC cells to hypoxic conditions (Wang et al. 2017; Tamukong et al. 2022), VEGFA is a key factor for new blood vessel formations, essential for tumor metastasis, and LOX contributes to ccRCC progression by increasing the stiffness of the collagen matrix, which in turn, facilitates the cellular migration (Di Stefano et al. 2016).

We explored the number of overlapping transcripts to understand the inter-tumor heterogeneity of alternative splicing between patients. As the PacBio Iso-Seq pipeline assigns transcript IDs randomly, we matched the transcripts based on their exon-boundaries as described before by Healey et al. (Healey et al. 2022). Using the Tama tool, we could detect 11,283 common transcripts from 4,746 genes after applying stringent filtering in every sample (see Methods). 2,393 transcripts were found only in ccRCC2, ccRCC4, and ccRCC5 PDOs, not in the Normal and ccRCC3 (Fig 4A). Transcripts that were not unique to a sample were found to be expressed in more cells (Figure 4B, Wilcoxon test, p-value < 2.2e-16). A comparison of the number of all matched transcripts revealed the highest similarities between Normal:ccRCC5 (Jaccard similarity index: 0.23) and ccRCC2:ccRCC5 PDOs (Jaccard similarity index: 0.23) followed by Normal:ccRCC2 (Jaccard similarity index: 0.22) (Supplementary Figure 5).

**Fig 4:**
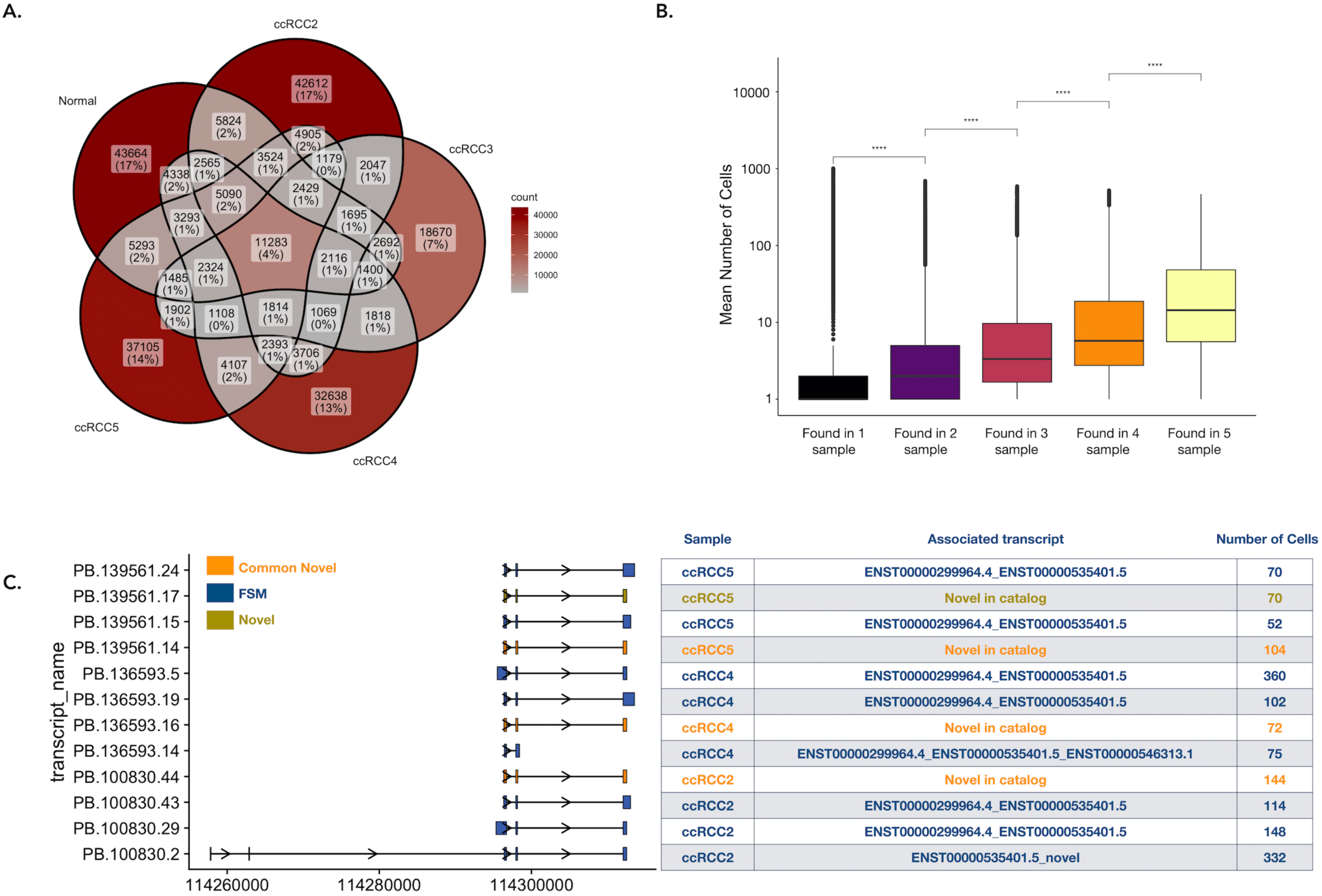
Shared transcripts across samples: (A). The number of overlapping transcripts across all samples. **(B).** Box plots showing the mean number of cells a transcript was found in a sample on average (Wilcoxon test, p<2.2e-16). **(C).** Top four transcripts of NNMT transcripts based on the number of cells across ccRCC2, ccRCC4, and ccRCC5 and their exon structures (left panel). The commonly found novel transcripts from ccRCC2, ccRCC4, and ccRCC5 are depicted in orange, FSM transcripts are shown in blue, a distinct novel transcript from ccRCC4 is highlighted in yellow. The table next to the transcript structures lists the SQANTI3 categories, aligned reference transcripts, and the number of cells in which the transcripts were identified (right panel).

The comparison of transcripts found explicitly in CA9+ or CA9- cells in each sample revealed 1,364 transcripts commonly detected in ccRCC cells of ccRCC2 and ccRCC5 PDOs, 48 transcripts between ccRCC4 and ccRCC5, and 100 between ccRCC2 and ccRCC4 (Supplementary Figure 6A). Next, we explored the splicing diversity between ccRCC and non-ccRCC cells. Interestingly, no preference was found for the number of novel transcripts in ccRCC and non-ccRCC cells (Fig. 3D).

. Explicitly expressed transcripts in ccRCC and non-ccRCC cells showed a diverse structural category pattern (Supplementary Figure 6B). Nevertheless, we observed 748 novel transcripts from 582 genes commonly found in ccRCC cells (Figure 3E, see Supplementary File 2 for a full list that were mostly associated with ccRCC-relevant pathways, including hypoxia signaling, glycolysis and oxidative phosphorylation (Fig. 3F). For example, the genes expressing highest number of common novel isoforms in CA9 expressing cells in the ccRCC5 and ccRCC2 were *NDUFA4L2* gene with 17 novel transcripts, and ANGPTL4 with 11 novel transcripts. Both genes play a role in ccRCC progression, as mentioned earlier.

One of the most frequently found novel transcripts belonged to the Nicotinamide N- Methyltransferase (NNMT) gene. It was categorized as NIC having a combination of known junctions between three exons (Fig. 4C).

### PCR validation experiments

We validated the novel transcript of NNMT using PCR (see Fig. 4C). The novel transcript of *NNMT* differed from the commonly found FSM isoform by the presence of 36 nucleotides at the end of the exon 2 (Fig. 5A). ORF prediction showed that the candidate novel isoform contained a stop codon at the beginning of the unique sequence. The novel isoform of NNMT has 121 amino acids, corresponding to part of the catalytic domain, NNMT_PNMT_TEMT compared to the canonical NNMT protein sequence (PDB ID: 3ROD, (Peng et al. 2011)) (Fig. 5B). Thus, the novel transcript lacks the complete binding pocket and likely any enzymatic activity. We successfully detected the unique region of the novel transcript using PCR and Sanger sequencing (Fig. 5D, 5E, NNMT Novel; Supplementary Fig. 7B). Additionally, we were able to detect the unique region of the FSM (NNMT Canonical). It should be noted that there is a known isoform of NNMT in the Ensembl database, ENST00000545255, which contains only two exons and shares 100% sequence similarity with the identified region of novel isoform in Sanger sequencing. However, the ENST00000545255 isoform was not identified in our data, which supports the conclusion that the identified isoform is indeed novel. Note that the validation of a novel transcript of TMEM91 gave non-conclusive results (see Supplementary Materials).

**Fig 5.**
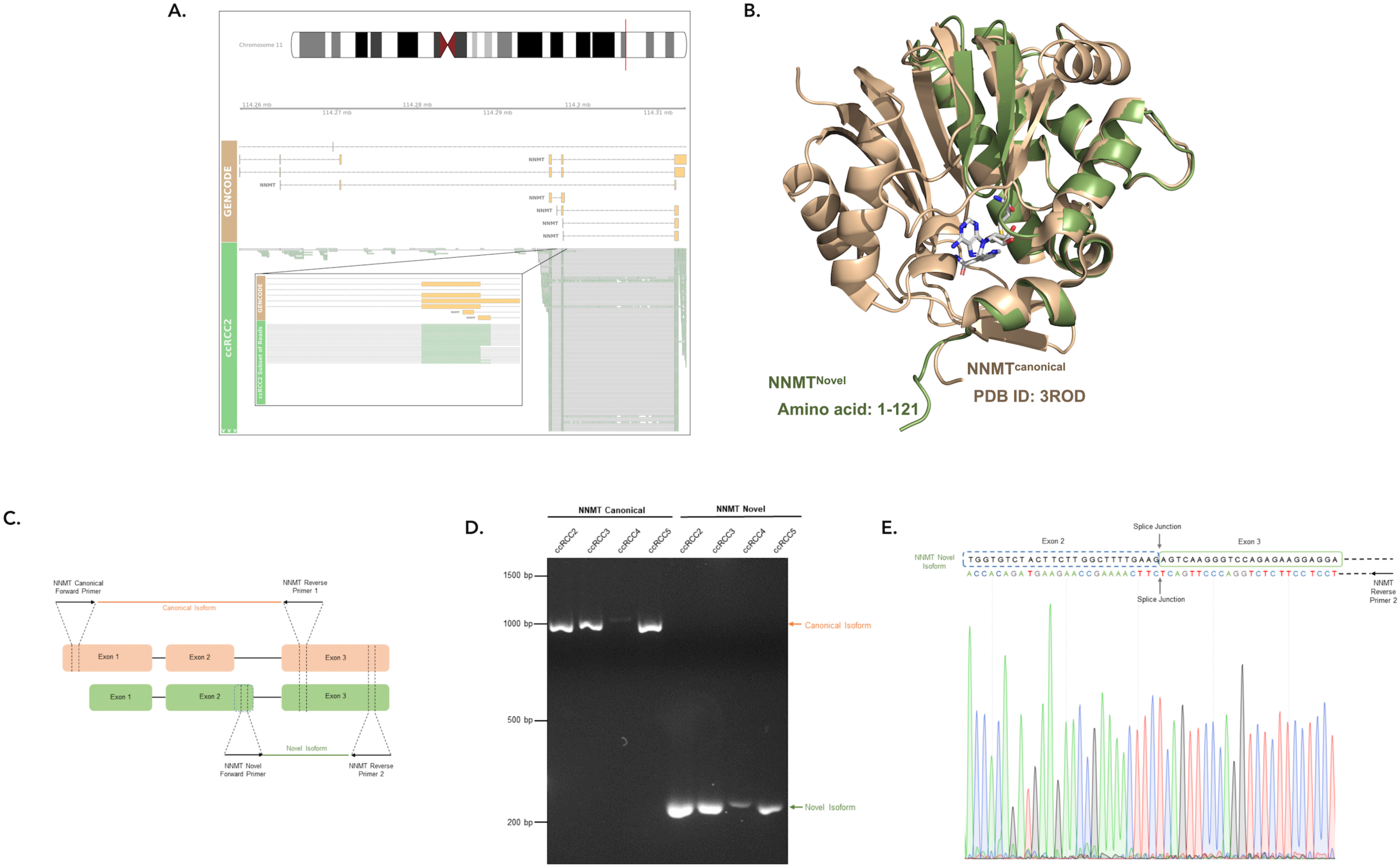
NNMT novel transcripts: **(A)** Reads that align to the NNMT novel transcript. **(B)** Alphafold3 structure of the NNMT novel isoform aligned to the canonical isoform of NNMT (PDB ID: 3ROD). **(C).** Representation of the primer design strategy to validate and sequence the novel and canonical transcripts of NNMT. In the novel isoform the dotted box indicates the position of the unique sequence at 3’ end of exon 2. **(D)**. Agarose gel (2%) electrophoresis image of PCR validation of different NNMT transcripts. All lanes are marked with corresponding tumor samples and product name. The canonical transcript is amplified with canonical forward primer and reverse primer 1 (see orange arrow). The novel transcript is amplified with a novel forward primer and reverse primer 2 (see green arrow). **(E).** Sanger sequencing result of NNMT novel transcript. Reverse sequencing confirmed the unique sequence at the 3’ end of exon 2 of the novel transcript. The splice junction between exon 2 and exon 3 is marked.

### Most Dominant Transcripts Switching Events in ccRCC Cells

As alternatively spliced transcripts can have different exons, they may result in different protein domains, disrupt protein interactions, or form interaction with new protein partners. Previous research has shown that most protein-coding genes have one most dominant transcript (MDT) expressed at a significantly higher level than any other transcript of the same gene. These dominant transcripts can be tissue-specific (Ezkurdia et al. 2015; Gonzàlez-Porta et al. 2013; Tung et al. 2022). We previously demonstrated that these MDTs switch during malignant transition in cancer, including in ccRCC (Kahraman et al. 2020). To explore variations in MDT profiles between ccRCC and non- ccRCC cells, we analyzed MDT distribution and their switches between ccRCC and non-ccRCC cells across three PDOs, ccRCC2, ccRCC4, and ccRCC5. The highest number of genes having MDT was found for ccRCC5 non-ccRCC cells (Fig. 6A). In total, we identified 7,986 unique cancer- specific MDTs in 571 single cells, ranging between one and 48switches per cell (Fig. 5A). Most of these switches were found only in one cell in a sample (78.1% in ccRCC2, 99.6% in ccRCC4, 88.4% in ccRCC5). Interestingly, no cancer-specific MDT was found in three samples (Fig. 6C, left panel). However,73 genes expressed different cMDT across all CA9+ samples (Fig. 6D, right panel), while 764 genes showed a cMDT in at least two ccRCC samples. Over-representation analysis of these genes revealed functional roles in RNA and mRNA splicing pathways, ubiquitin dependent protein catabolic process, regulation of mRNA metabolic process, and mitochondrial translation (Fig. 6D). The most frequently found cancer-specific MDT that was expressed in 115 cells of ccRCC2 was APH1A (Fig. 6E). The APH1A gene encodes for the transmembrane protein Aph-1. This protein is a part of the gamma-secretase complex, having a role in the cleavage of various transmembrane proteins, including proteins associated with cancer, such as Notch, ErbB4, CD44, VEGFR, etc (Song et al. 2023). The ccRCC MDT aligns to ENST00000369109.8 with an alternative 5’ end. It had seven exons and encoded 265 amino acid-long protein. The non-ccRCC MDT mapped to ENST00000360244.8 with an alternative 5’ end consisting of six exons and encoding for 247 amino acid long protein (Fig. 6E). On GTEx, both isoforms were found to be expressed in high abundance; ENST00000369109.8 is the most abundant isoform on Kidney Medulla, while the ENST00000360244.8 is the most abundant transcript in Kidney Cortex (https://www.gtexportal.org). The expression of both isoforms in the cells expressing ENST00000369109 as cMDT and in the normal cells are shown in Fig. 6F.

**Fig 6:**
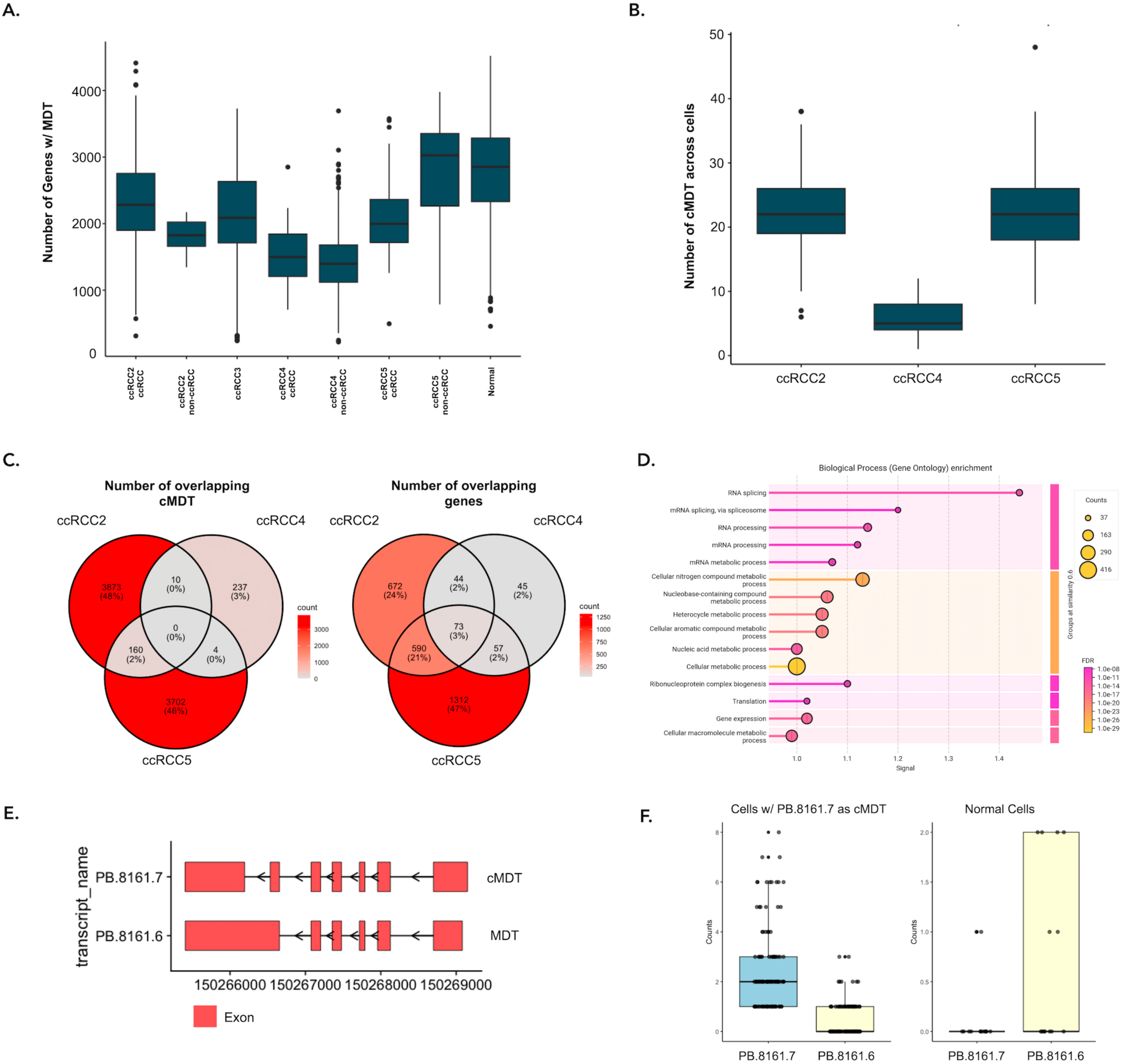
MDTs and MDT Switches between ccRCC and non-ccRCC cells: (A). Distribution of the number of MDTs in PDOs. **(B).** Distribution of the number of cMDT in ccRCC2, ccRCC4, and ccRCC5 PDOs. **(C).** The number of overlapping isoforms (left-panel) and genes (right-panel) showing transcript switching events across three datasets. **(D).** GO term enrichment analysis of genes commonly showing transcript switching events in any of ccRCC2 ccRCC5, and ccRCC4 PDO datasets. **(E).** Exon structures of ccRCC (PB.8161.7) and non-ccRCC (PB.8161.6) MDTs. **(F).** UMI counts of cMDT and MDT APH1A transcripts in ccRCC and non- ccRCC cells.

In ccRCC5 PDO cells, the most frequently found cMDT belonged to the gene TMEM161B divergent transcript (*TMEM161B-DT)* which is a long noncoding RNA. Higher expression of TMEM161B-DT has been associated with malignancy of glioma cells (Chen et al. 2021) while it was found to be downregulated in oesophageal squamous cell carcinoma (Shi et al. 2021)Here, we identified a cMDT of TMEM161B-DT in 71 ccRCC cells of ccRCC5 PDO (Supplementary File 3). The cMDT had three exons, mapped to ENST00000665319.2 with an alternative 3’ end. In non-ccRCC cells of ccRCC5, we identified diverse MDTs classified as FSM, NIC, or ISM, having three to four exons.

A differential splicing analysis with edgeR using *acorde* (Arzalluz-Luque et al. 2022) revealed three upregulated transcripts in non-ccRCC cells within the ccRCC2 sample that matched MDTs in the Normal sample (Supplementary Table 3).

## Discussion

The recent advent of single-cell long-read sequencing technologies provides a unique opportunity to gain insight into intra- and inter-tumor heterogeneity of tumors and to discover potential novel predictive biomarkers. To reveal the heterogeneity in ccRCC, we utilised the MAS-Seq single-cell long-read sequencing protocol of PacBio. We generated a comprehensive catalogue of known and novel transcripts for one normal and four ccRCC Patient-Derived Organoids (PDOs) without employing short-read single-cell sequencing data. PDOs with the highest number of sequenced cells, had, as expected, the least number of detected transcripts per gene per cell. However, sequencing a low number of cells might also cause a loss of essential cell diversity in the samples.

Here, we uncovered over 256,088 unique isoforms across all samples, of which 114,434 (44.7%) are novel transcripts with new combinations of known exons or new junctions. To interpret the biological impact of transcripts sequenced, we investigated the prevalence across cells together with their protein-coding capability. Our analysis revealed that, as expected, conserved well- characterized transcripts were more widely expressed across all cells and samples. In contrast, on average 61% of identified transcripts were found only in one cell suggesting rare diversity that would need further investigation with a higher number of cells. Frequently identified known and novel transcripts had more complete open reading frames, emphasizing their protein-coding capability.

As expected, the highest proportion of transcripts in our data was found to be incomplete splice matches. These transcripts showed the least fraction of complete ORFs and the highest disordered score for complete ORFs. Proteins encoded by these transcripts may exhibit enhanced functional diversity or regulatory capacity due to the lack of a stable protein structure. To understand the splicing diversity between ccRCC and non-ccRCC cells, we investigated explicitly expressed transcripts in each category. ccRCC cells tended to have unique novel transcripts in ccRCC-related pathways (e.g. for oxidative phosphorylation, hypoxia, and glycolysis), proposing a contribution to ccRCC cancer progression. Our most dominant switch analysis between ccRCC and non-ccRCC cells revealedmany cell and sample-specific switching events. Nevertheless, genes showing switching events were often part of the mRNA-splicing pathway, highlighting a pivotal role alternative splicing regulation. But abundant mRNA transcripts must not necessarily translate to high abundant proteins. Miller et al. demonstrated how long-read data can drive the validation process of new protein isoforms. For their validation, the authors constructed a protein reference database with full-length transcript sequences in order to use the database for querying the mass-spectrometry-based proteomics data. The authors were able to confirm novel peptide and translated intronic sequences. The total number of these identifications was low but highlighted the possibility of transcript translations commonly ignored or overseen in classical proteomics experiments (Miller et al. 2022).

Our study provides an insight into the complex and under-explored functional diversity of cells in ccRCC. In our data, where possible, we have meticulously addressed the issue of potential artifacts and biases potentially introduced by sample processing or data analysis., Despite our efforts to minimize any artifacts some limitations might still have remained. One issue we could not address or quantify is the introduction of artifacts in the PCR amplification, an essential step in the MAS-ISO-seq library protocol. However, a recent study by Lee *et al*. demonstrated a good overlap of transcript abundances assessed with PCR amplified cDNA molecules and direct RNA sequencing using Oxford Nanopore sequencing (Lee et al. 2023). Another issue could be associated with the difficulty in delineating the actual isoform architecture disguised by any transcript degradation, fragmentation, or incompleteness. Iso-Seq addresses the issue by flexibly merging isoforms with differing internal and external junctions. However, the parameters might not be optimized to cell, sample, or tissue types. Lastly, Iso-Seq works on a per-sample basis and provides arbitrary isoform IDs which cannot be matched between samples. The tool Tama merge, that was used in this work, does not take into account the sequence identity between matched transcripts, across samples, which can mask some of the isoforms’ diversity. Existent tools that can perform multi-sample isoform discovery and quantification, including bambu (Chen et al. 2023), employ different algorithms (e.g., machine-learning based) that often produce different sets of transcripts. This raises questions about the isoform collapsing parameters, read correction methods, and the sufficient amount of evidence that is required to call a transcript novel. We think the issue can be addressed with an investigation of wider depth of tissue types and with the use of molecular validation assays.

In conclusion, single-cell long-read sequencing of patient-derived organoids offers an unprecedented detailed view of the transcriptome landscape of individual cancer patients. It reveals hundreds of thousands of novel transcripts, of which only the minority are commonly expressed in single and multiple patients, highlighting the intra- and inter-tumor heterogeneity of ccRCC. The discovery of frequently found novel transcripts provides insights into cancer progression and a new avenue for discovering potential novel biomarkers or therapeutic targets. The functional role of the commonly expressed novel transcripts remains to be further explored and validated.

## Methods

### Generation and Characterization of ccRCC Patient-Derived Organoid Samples

Patient tissue samples were provided by the Department of Pathology and Molecular Pathology at University Hospital Zürich. The tissues were collected and biobanked according to previously described procedures (Bolck et al. 2019). The study was approved by the local Ethics Committee (BASEC# 201 9-01 959) and in agreement with the Swiss Human Research Act (Swiss Human Research Act). All patients gave written consent. Organoids were established as previously described (Bolck et al. 2021). Surgically resected renal tissue was reviewed by a pathologist with specialization in uropathology (Holger Moch) and suitable specimens were stored at 4 °C in transport media (RPMI (Gibco) with 10 % fetal calf serum (FCS, Gibco) and Antibiotic-Antimycotic® (Gibco)). For organoid derivation, tissue specimens were processed within 24 hours by rinsing them once with PBS followed by finely cutting and digesting them in 0.025 mg/ml Liberase (Roche) for 15 min at 37°C. The slurry was passed through a 100 µm cell strainer and centrifuged at 1000 rpm for 5 min. Cells were washed once with PBS and erythrocytes were lysed in ACK buffer (150 mM NH_4_Cl, 10 mM KHCO_3_, 100 mM EDTA) for 2 min at room temperature. After a final wash with PBS, appropriate amounts of cell suspension were resuspended in CK3D medium (Advanced DMEM/F12 (Gibco) with

- 1X Glutamax (Gibco)
- 10 mM HEPES (Sigma-Aldrich)
- 1.5X B27 supplement (Gibco)
- Antibiotic-Antimycotic (Gibco)
- 1 mM N-Acetylcysteine (Sigma-Aldrich)
- 50 ng/mL Human Recombinant EGF (Sigma-Aldrich)
- 100 ng/mL Human Recombinant FGF-10 (Peprotech)
- 1 mM A-83-01 (Sigma-Aldrich)
- 10 mM Nicotinamide (Sigma-Aldrich)
- 100 nM Hydrocortisone (HC, Sigma-Aldrich)
- 0.5 mg/ml epinephrine (Sigma-Aldrich)
- 4 pg/mL Triiodo-L-thyronine (T3, Promocell)
- R-Spondin (conditioned media, self-made)

The composition was mixed with two volumes of growth factor reduced Matrigel (Corning). Drops of cell suspension/Matrigel were distributed in a 6-well low attachment cell culture plate (Sarstedt) and allowed to solidify for 30 min at 37 °C, upon which CK3D media was added to cover the drops. To evaluate the growth of PDOs, bright-field images were captured using a microscope. Organoids at approximately 100-500 um were passaged, and at least 10,000 cells were collected for cell model validation using targeted DNA sequencing of the *VHL* gene. To achieve this, DNA was isolated using the Maxwell® 16 DNA Purification Kit (Promega) and corresponding Maxwell instrument. PCR and sequencing of *VHL* were performed as previously described (Rechsteiner et al. 2011).

### Full-length single-cell isoform sequencing and data processing of PDO cells via MAS-ISO-Seq

To obtain single cell suspension, cell culture media was removed and PDOs from one well of a ULA 6-well plate were collected in ice-cold Cell Recovery Solution (Corning) and incubated for 1 hour at 4°C to resolve the Matrigel. Subsequently, PDOs were dissociated with TrypLE by incubation on a thermal shaker set to 37 °C, 300 rpm. Every 2 min, the samples were picked up and mechanically dissociated by pipetting up and down and the progress of dissociation was evaluated under a microscope using a small fraction of the cells and tryphan blue. After dissociation, PBS supplemented with 20 % FBS, was added to stop the reaction. Samples were centrifuged at 1000 *g* for 5 min and the supernatant was aspirated. The pellet was washed once in 1X PBS with 0.04 % BSA and filtered through a 70 µm strainer. Finally, cells were counted and diluted to the target cell concentration using PBS with 0.04 % BSA. Cell viability and concentration were determined using a LUNA-FX7 Automated Cell Counter (Logos).

### Generation of full-length cDNA with 10x Genomics platform and PacBio MAS-Seq library preparation and sequencing

10x Genomics Chromium platform was used to analyze the dissociated organoid cells (Zheng et al. 2017). We targeted to recover 700 cells per library preparation to have a greater sequencing depth using the PacBio platform. Library preparation was conducted following the 10x Genomics Single Cell 3’ Reagent Kits v3.1 (Dual Index) User Guide. Cells were combined with a master mix containing reverse transcription reagents. The single-cell 3’ v3.1 gel beads, which carry the Illumina TruSeq Read1, a 16bp 10x barcode, a 12bp UMI, and a poly-dT primer, were loaded onto the chip along with oil for the emulsion reaction. Chromium X partitioned the cells into nanoliter-scale gel beads in emulsion (GEMs) followed by reverse transcription. All cDNAs within a GEM, representing one cell, shared a common barcode. After the reverse transcription reaction, the GEMs were broken, and the full-length cDNAs were captured by MyOne SILANE Dynabeads and then amplified. The amplified cDNA underwent cleanup with SPRI beads, followed by qualitative and quantitative analysis using an Agilent 4200 TapeStation High Sensitivity D5000 ScreenTape and Qubit 1X dsDNA High Sensitivity Kit (Thermo Fisher Scientific). The single-cell full-length cDNAs were directed for single-cell MAS-Seq (Multiplexed Arrays Sequencing) library preparation using the MAS-Seq 10x Single Cell 3’ kit (Pacific Bioscience, CA, USA). Template switch oligo (TSO) priming artifacts generated during 10x cDNA synthesis were removed in the PCR step with a modified PCR primer (MAS capture primer Fwd) to incorporate a biotin tag into desired cDNA products followed by capture with streptavidin-coated MAS beads. TSO artefact-free cDNA was then further directed for the incorporation of programmable segmentation adapter sequences in 16 parallel PCR reactions/sample followed by directional assembly of amplified cDNA segments into a linear array. The obtained 10-15 kb fragments were subjected to DNA damage repair and nuclease treatment. The quality and quantity of the single-cell MAS-Seq libraries were assessed with Qubit 1X dsDNA High Sensitivity Kit (Thermo Fisher Scientific) and pulse-field capillary electrophoresis system Femto Pulse (Agilent), respectively. Each single-cell MAS-seq library was used to prepare the sequencing DNA-Polymerase complex using 3.2 binding chemistry and further sequenced on a single 8M SMRT cell (Pacific Bioscience), on Sequel IIe sequencer (Pacific Bioscience) yielding in ∼ 2 M HiFi reads and∼ 30M segmented reads per sample.

### Short Read sequencing

The second part of the cDNA was used for Illumina sequencing library preparation, following the 10x Genomics Chromium Single Cell 3’ Reagent (v3.1 Chemistry Dual Index) protocol as described above. The cDNA was enzymatically sheared to a target size of 200-300 bp, and Illumina sequencing libraries were constructed. This process included end repair and A-tailing, adapter ligation, a sample index PCR, and SPRI bead clean-ups with double-sided size selection. The sample index PCR added a unique dual index for sample multiplexing during sequencing. The final libraries contained P5 and P7 primers used in Illumina bridge amplification. Sequencing was performed using paired-end 28-91 bp sequencing on an Illumina Novaseq 6000 to achieve approximately 300,000 reads per cell.

### SMRTLink Iso-Seq pipeline

In our study, we utilized the "Read Segmentation and Iso-Seq workflow" from SMRTLink version 11.1 to process our long-read sequencing data. For two specific samples, Normal and ccRCC2, we combined the data from three SMRTcells to enhance coverage. Within the pipeline, the HiFi reads were converted into segmented reads using the skera tool, followed by processing with the Iso-Seq for removal of cDNA primers and barcode and UMI tags, reorientation, trimming of poly-A tails, cell barcode correction, real cell identification and PCR deduplication via clustering by UMI and cell barcodes. The reads were then aligned to the human genome (GRCh38.p13) using pbmm2. We verified the presence of the selected 10x cell barcodes using the GenomicAlignments R package (Lawrence et al. 2013).

### Full-length single-cell data analysis

#### Isoform Filtering

After mapping, isoforms were collapsed into a unique set of transcripts with Iso-Seq using the default options, setting –max-fuzzy-junction to 5bp, –max-5p-diff to 1000bp, –max-3p-diff to 100bp, –min- aln-coverage to 0.99, –min-aln-identity to 0.95, –max-batch-mem 4096, and –split-group-size to 100. In addition, at the Isoseq collapse step, reads that mapped chimerically or mapped with low identity were filtered out. The *pigeon make-seurat* function was run on the remaining reads to generate the gene count matrices. Subsequently, pigeon was used to classify the unique isoforms into SQANTI3 classification categories (Pardo-Palacios et al. 2023). After isoform classification, pigeon filtered out intra-priming (with accidental priming of adenine stretches in the genomic position downstream of the 3’ end), RT switching (reverse transcriptase template switching) and low coverage/non-canonical isoforms (having non-canonical splice junctions).

In addition to the pigeon-based filtering, we manually filtered transcripts based on their Transcription Start Site (TSS) ratio, their distance to the gene’s TSS and Transcription Termination Site (TTS) and their distance to the gene’s CAGE peak. We calculated the TSS ratio using Illumina short reads as an input to SQANTI3’s stand-alone sqanti3_qc.py function and discarded any 5’ end-degraded transcripts. We used different filtering criteria for each SQANTI category: FSM isoforms: TTS ratio > 1; ISM, NNC, NIC, and other isoforms: TTS ratio > 1, distance to CAGE peak ≤ 50 bp, and distance to the gene’s TTS and TSS ≤ 50 bp. All other isoforms were discarded for downstream analysis.

The isoform count matrices were generated with the *pigeon make-seurat* function on the filtered isoforms with default parameters. Reads mapping to mitochondrial and ribosomal genes were not retained during isoform and gene count matrix generation. Additionally, we only kept the cells with mitochondrial content <30% for the downstream analysis.

#### Transcript Types and Their Prevalence Across Cells

We calculated the percentage of structural categories and their length in each sample using filtered scisoseq_classification.filtered_lite_classification.txt files. We then checked transcript prevalence across varying cell number ranges, and the number of transcripts per gene and cell.

#### Functional Annotation of Long-read Sequencing Transcripts

Open Reading Frames (ORFs) were identified on long-read transcript sequences listed in fasta files from the Iso-Seq collapse function using Transdecoder v5.7.1 (Haas BJ.). The Transdecoder.LongOrfs function was used to predict all possible ORFs with a length ≥100 nucleotides. To calculate protein sequences from the predicted ORFs, an extensive human reference database containing 226,259 canonical and alternatively spliced isoform protein sequences was generated using Uniprot (release date: 2023-11). The predicted ORFs were aligned to this database via blastp, setting the e-value to 1e-5. In addition, hmmscan v3.4 was applied to predict potential Pfam domains using the Pfam database (release date: 2023-09-12) with a maximum e-value of 1e-10. The results from both hmmscan and blastp were used to predict the final ORFs using the Transdecoder.Predict function. We then selected one ORF for each transcript based on the highest score assigned by TransDecoder. We applied iupred2a on the transcripts having complete ORFs to predict their intrinsically disordered regions (IDRs). A residue was annotated as ordered or disordered, if its iupred2a score was below or above 0.5, respectively. We calculated the percentage of disordered residues for each transcript and assigned a percentage disordered score for each transcript.

#### Transcript Matching among Samples

Due to Iso-seq assigning transcript IDs randomly, we first converted all sqanti_classification.filtered_lite.gff files to BED format using bedparse’s gtf2bed function (Healey et al. 2022). A “geneID;TranscriptID” column was added to the BED file. Tama’s tama_merge.py function was used to combine all transcript ids among samples using their exon and junction coordinates. Mismatches up to 50 and 100 nucleotide from the 5’ and 3’ ends, respectively, were accepted, as well as mismatches 5 nucleotides from any exon junction. The similarities of the samples were calculated in R using the Jaccard similarity matrix, i.e. the number of overlapping transcript IDs divided by the total number of transcripts found in two samples. The heatmaps were visualized using the pheatmap function in R, and the number of overlapping transcripts was plotted by UpsetR’s upset function (Conway et al. 2017).

#### Cell Type Annotations

Seurat (Hao et al. 2024) was used for quality control and integration of the samples using the output files of the Iso-Seq make-seurat function. For gene-level analysis, each sample was normalized by the SCTransform function. 3000 features were selected using SelectIntegrationFeatures, and anchors for integration were identified with FindIntegrationAnchors. The samples were integrated with the IntegrateData function using the SCT normalization. Subsequently, the PCA, and UMAP analyses were performed using the RunPCA and RunUMAP functions, respectively. Markers for each cluster were defined with the PrepSCTFindMarkers and FindAllMarkers functions. To categorize the cells in each PDO, we analyzed the samples separately. SCT normalized gene expression matrices were scaled, and the cells were categorized into two categories using the scGate R package (Andreatta et al. 2022) by defining the CA9 as a ccRCC positive marker. The other cells were assigned as non-ccRCC. We used SCpubr R package to visualize marker expressions and clusters (Blanco-Carmona 2022). Genes expressing ccRCC-specific novel transcripts in ccRCC cells of ccRCC2,ccRCC4 and ccRCC5 were analyzed using ClusterProfiler’s enricher function (Yu et al. 2012) For the analysis a hallmark gene set from MsigDB was used as the background gene set (Liberzon et al. 2015). An overrepresentation analysis was performed setting pValueCutoff = 0.05, qvalueCutoff = 0.1, and pAdjustMethod = BH (Benjamini-Hochberg). In addition, we annotated cells using manual curation of ccRCC and kidney related markers using sc- Type (Ianevski et al. 2022). We used the following markers for the annotation of cells:

- ccRCC cells: CA9, ANGPTL4, NDUFA4L2, LOX, VEGFA, VIM, and EGLN3.
- Proximal Tubule Cells (PTC); EPCAM, PAX8, GGT1, and RIDA.
- Stromal Cells: ACTA2, FAP, COL1A1, and COL1A2.
- Endothelial Vascular Cells: CDH5, FLT1, PECAM1, and KDR.
- Immune Cells: CD3, CD8A, PD1, CTLA4, CD68, CD163, and CD11C.
- Stem Cells: ALDH1A1, SOX2, and CD44.
- Mesenchymal Cell: VIM, FN1, SNAI1, SNAI2, ZEB1, and ZEB2.
- Epithelial-mesenchymal transition (EMT): CDH2, TWIST1, MMP2, and MMP9.

#### Most Dominant Transcripts Switches between ccRCC and non-ccRCC cells

To assess Most Dominant Transcripts (MDTs) and cancer-specific MDTs (cMDTs) in our 5 ccRCC samples, we have used transcript UMI counts in each sample. Each MDT was required to have at least two times higher UMI counts than the second most abundant transcript (Kahraman et al. 2020). Orphan transcript of genes were automatically counted as MDT. cMDT were computed based on the comparison of MDTs between ccRCC and non-ccRCC cells using following strict rules:

- cMDT are unique to ccRCC cells.
- For at least 20% of non-ccRCC cells, a distinct MDT of the same gene exists.
- If an MDT and potential cMDT mapped to the same transcripts within the sample, cMDT was discarded.
- UMI counts of MDTs in ccRCC cells should be higher than the mean of the MDTs’ UMI count in non-ccRCC cells.

A cMDT was identified when an MDT switch event fulfilled all criteria. STRING db was used for the enrichment analysis of genes showing MDT switches between ccRCC2 and ccRCC4, and ccRCC5 PDOs. For the enrichment analysis, the human gene list was used as a background (Szklarczyk et al. 2023). ggVennDiagram R package was utilized to generate a Venn diagram of overlapping cancer-specific MDTs among samples (Gao et al. 2021). Exon structures of the transcripts were generated with the ggtranscript R package (Gustavsson et al. 2022).

In addition, a differential isoform expression analysis was performed between ccRCC and non-ccRCC cells in each sample using the acorde software (Arzalluz-Luque et al. 2022). The software calculates cell-level weights for each isoform using ZinBWaVE R package (Risso et al. 2018) followed by performing differential expression with DESeq2 and edgeR.

#### Visualization of NNMT Reads and Structural Modelling

The protein structure of the novel NNMT isoform was modeled using AlphaFold3 (Abramson et al. 2024) based on its ORF sequence. The structure was rendered with PyMOL (Schrödinger, LLC.). Sequence reads were visualized using Gviz (Hahne and Ivanek 2016) and GenomicRanges (Lawrence et al. 2013).

### PCR Validations

To validate the isoforms using PCR, we targeted two novel isoforms of two genes, NNMT and TMEM91. We selected these novel isoforms based on their frequency and presence across samples. The novel isoform of NNMT was classified as Novel In Catalog (NIC) by SQANTI3. It was found in all samples (IDs: PB.100830.44 in ccRCC2, PB.139561.14 in ccRCC5, PB.136593.16 in ccRCC4, PB.130901.11 in Normal). The novel isoform of TMEM91 was classified as Novel not In Catalog (NNC) by SQANTI3. The transcript was identified predominantly in CA9 expressing ccRCC cells of ccRCC2 and ccRCC5 PDOs.

For the PCR experiment, total RNA was isolated directly from corresponding frozen tissue samples of ccRCC2, ccRCC3, ccRC4 and ccRRC5 using Maxwell RSC simplyRNA Tissue (promega, AS1340). 500 ng RNA was used to synthesize cDNA by qScript cDNA Synthesis Kit (Quanta bio, 95048-100) following the manufacturer’s protocol. Synthesized cDNA was used as a template for PCR amplification. In order to capture novel and canonical isoforms, we designed three types of primers against the novel transcripts:

- Common_primer: targeting sequences shared in both canonical and novel isoforms
- Canonical_primer: targeting sequences unique to canonical isoform
- Novel_primer: targeting sequences unique to novel isoform

For NNMT: A forward primer was specifically designed against the unique sequence of the novel isoform at the end of exon 2. To detect the canonical isoform, another forward primer was designed to span the unique sequence of the canonical transcript at exon 1. Both the reverse primers were designed against different regions of exon 3.

For TMEM91: A forward primer specific to the novel isoform was designed to span exon 1 of the novel transcript. Additionally, a forward primer was designed against the sequence shared between canonical and novel isoforms, corresponding to exon 2 in novel and exon 1 in canonical (mapping to ENST00000392002.7) transcripts. Reverse primer was designed to target a shared region of exon 3 (novel isoform)/exon 2 (canonical isoform).

All the primers were designed using Primer3 software (Untergasser et al. 2012) and synthesized by Microsynth AG. The details of primer sequences and primer pair combinations are listed in Supplementary Table 1 and 2, respectively. PCR amplification was performed using AmpliTaq Gold DNA Polymerase (Applied Biosystems, 4311806) following the manufacturer’s protocol. The amplified products were subjected to agarose gel electrophoresis (2%) and visualized with GelRed (Biotium, 41003-1). The PCR products were further purified by the MinElute PCR Purification Kit (Qiangen, 28006) and validated by Sanger sequencing at Microsynth AG.

## Data Access

The raw single-cell long-read RNA sequencing data generated in this study have been submitted to the European Nucleotide Archive (ENA: https://www.ebi.ac.uk/ena) under the accession number PRJEB73513.

The codes used in the manuscript can be found at: https://github.com/KarakulakTulay/ccRCC_scLongRead

## Funding

This work was funded by Krebsliga Zurich and the EMDO foundation

## Competing Interest Statement

None declared.

## Acknowledgments

We would like to thank Mark Robinson from UZH for valuable discussions about single-cell data analysis. We also would like to thank Harini Lakshminarayanan and Adriana Von Teichmann from the Department of Pathology and Molecular Pathology of USZ for their technical assistance during the preparation of PDO samples for the single-cell analysis.

## Notes

### Competing Interest Statement

The authors have declared no competing interest.

### Summary of Updates

We implemented stringent filtering criteria to enhance transcript reliability, as detailed in the newly added ‘Isoform Filtering’ section of the manuscript. The manuscript and all figures have been updated accordingly. Additionally, we validated one novel transcript experimentally using PCR and Sanger sequencing, with corresponding methods and results incorporated into the manuscript.

