## Supplementary Material for "Heterogeneous and Novel Transcript Expression in Single Cells of Patient-Derived ccRCC Organoids"

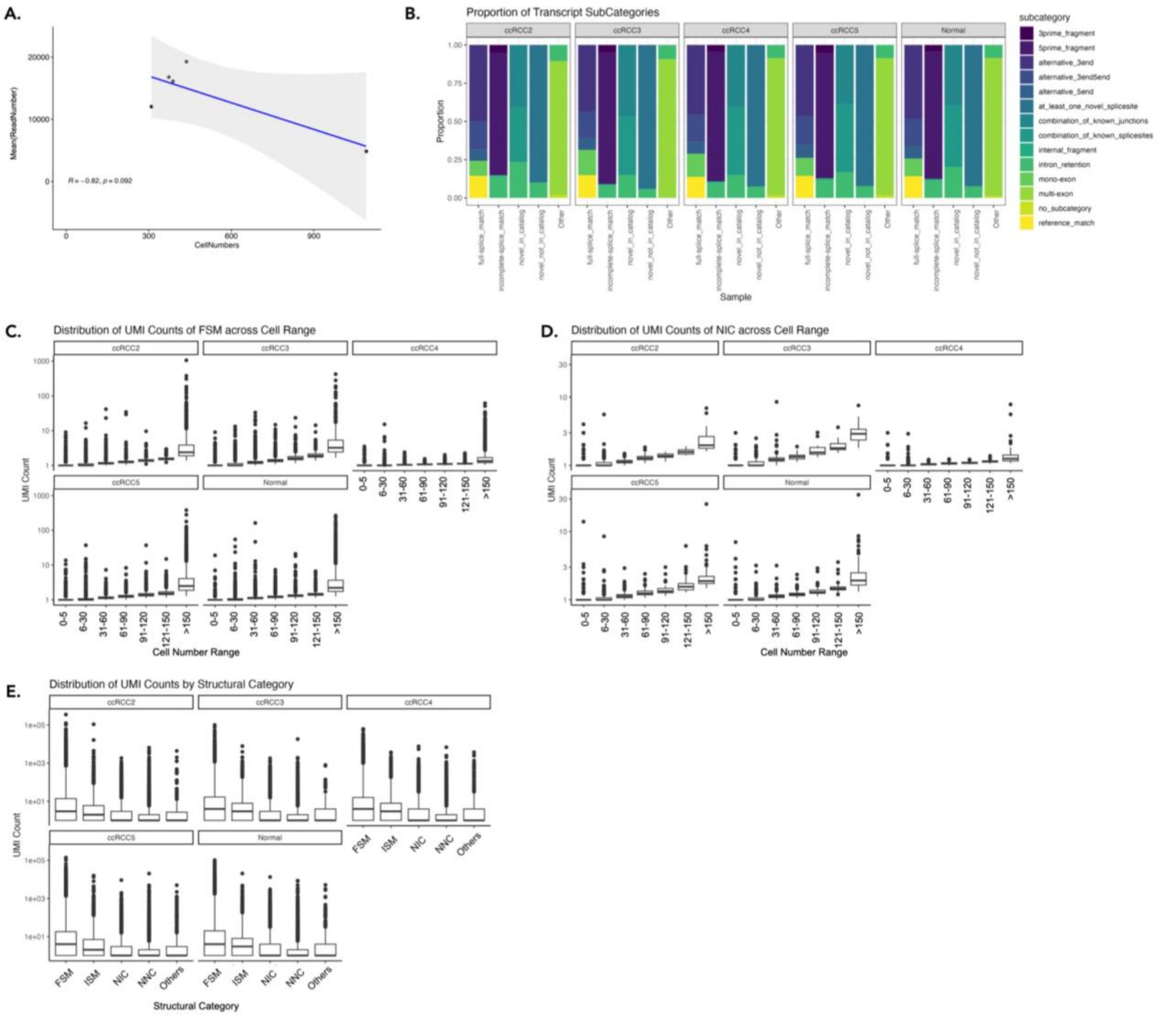


**Supplementary Figure 1: General statistics on transcripts found in three cells that express at least 100 genes/transcripts: (A).** Pearson Correlation between number of cells and mean of read number on the log2 scale (R=-0.81, p=0.096). **(B).** Proportions of Transcript Sub-categories Within Main Structural Categories for Each Sample. The x-axis represents the main structural categories as defined by SQANTI3 classification. Each bar graph depicts the fraction of the corresponding subcategories. **(C).** Distribution of UMI counts by cell nuof full-splice-match (FSM) transcripts across varying cell number range. **(D).** Distribution of UMI counts of Novel In Catalog transcripts (NIC) across varying cell number ranges. **(E).** Distribution of UMI counts across different structural categories of each sample. The Y-axis is on log10 scale in C, D, and normalised by the number of cell a transcript found in.

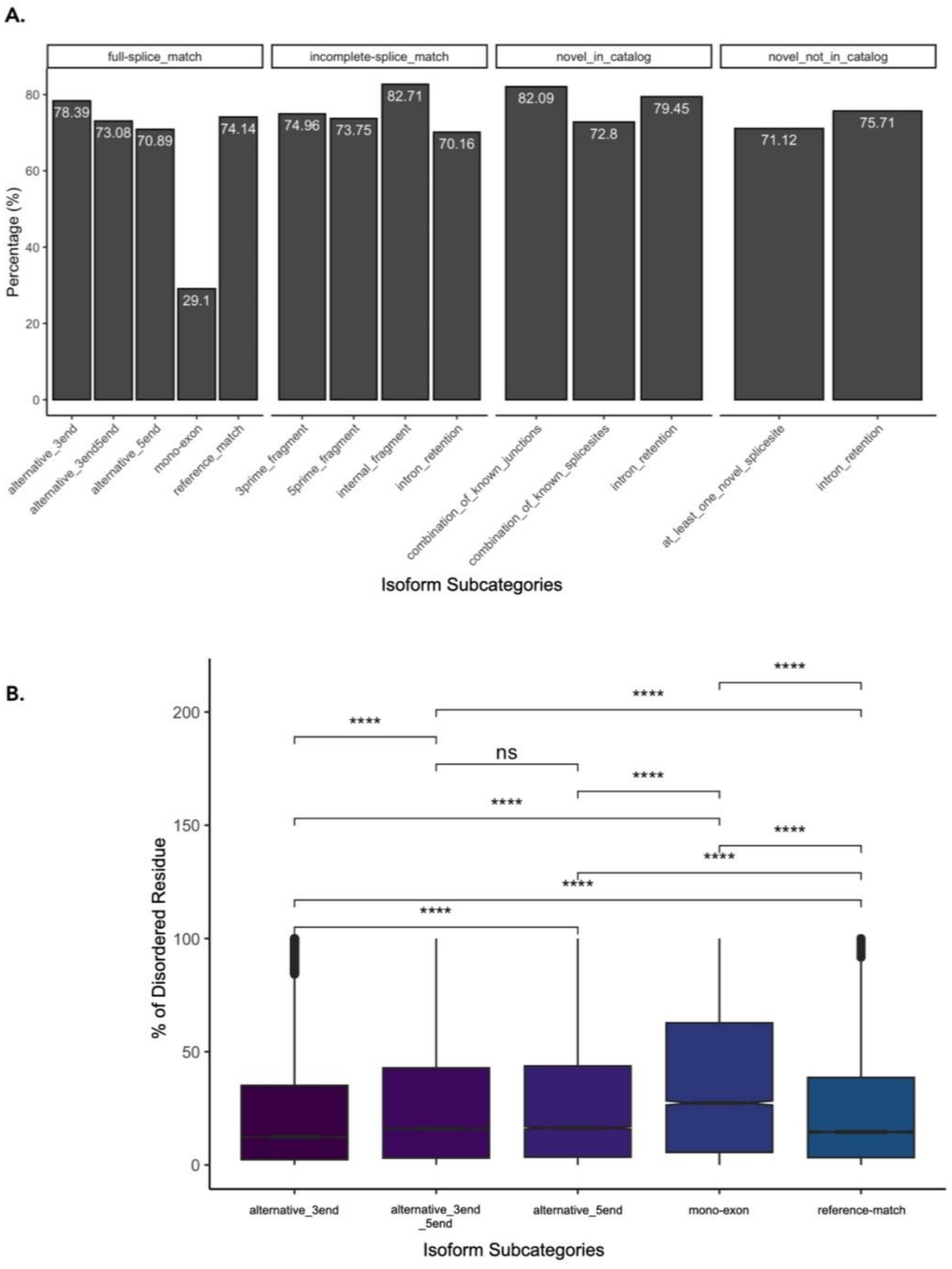


**Supplementary Figure 2: (A).** Percentage of ORF hit across isoform subcategories **(B).** Percentage of disordered regions across FSM subcategories.

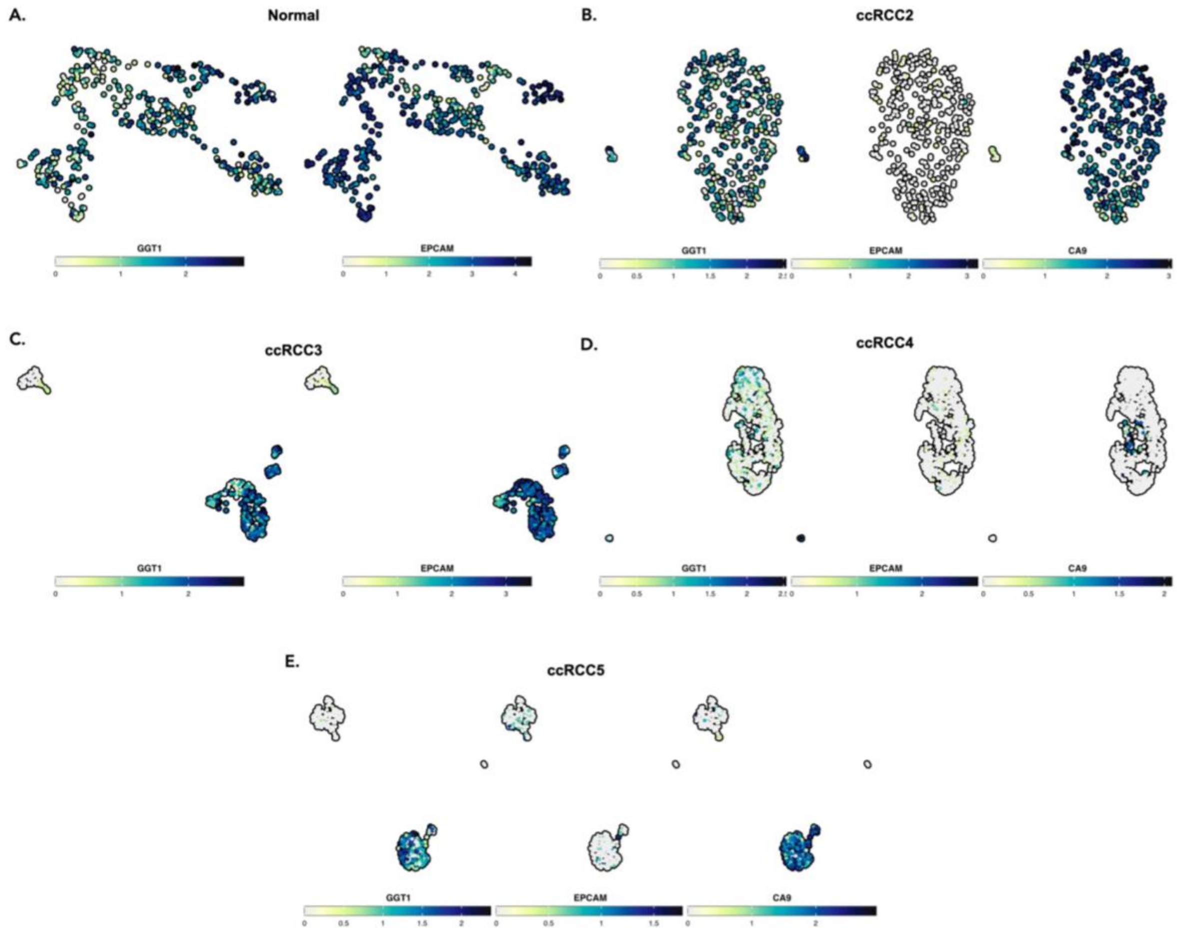


**Supplementary Figure 3:** Expression Pattern of GGT1, EPCAM, CA9 in (A) Normal PDO, (B) ccRCC2 PDO, (C) ccRCC3 PDO, (D) ccRCC4 PDO and (E) ccRCC5 PDO.

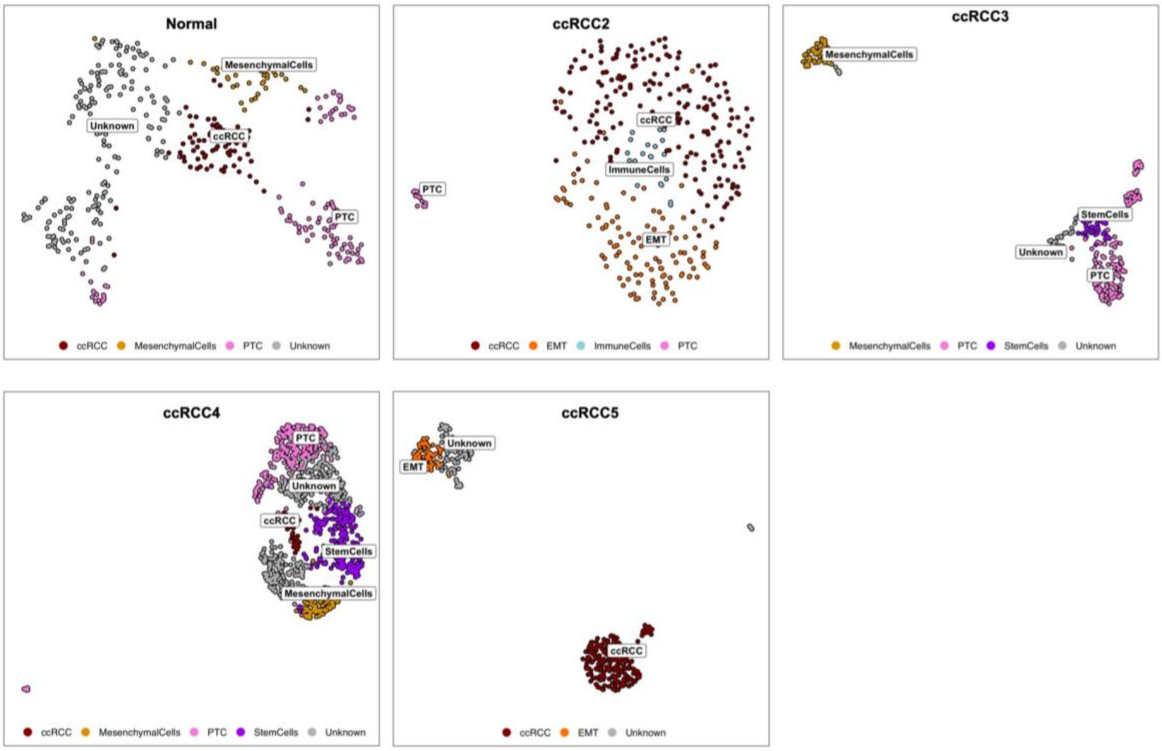


**Supplementary Figure 4:** Cell-type annotation of each sample with manually curated list of genes. PTC: Proximal tubule cells, EMT: Epithelial-mesenchymal transition.

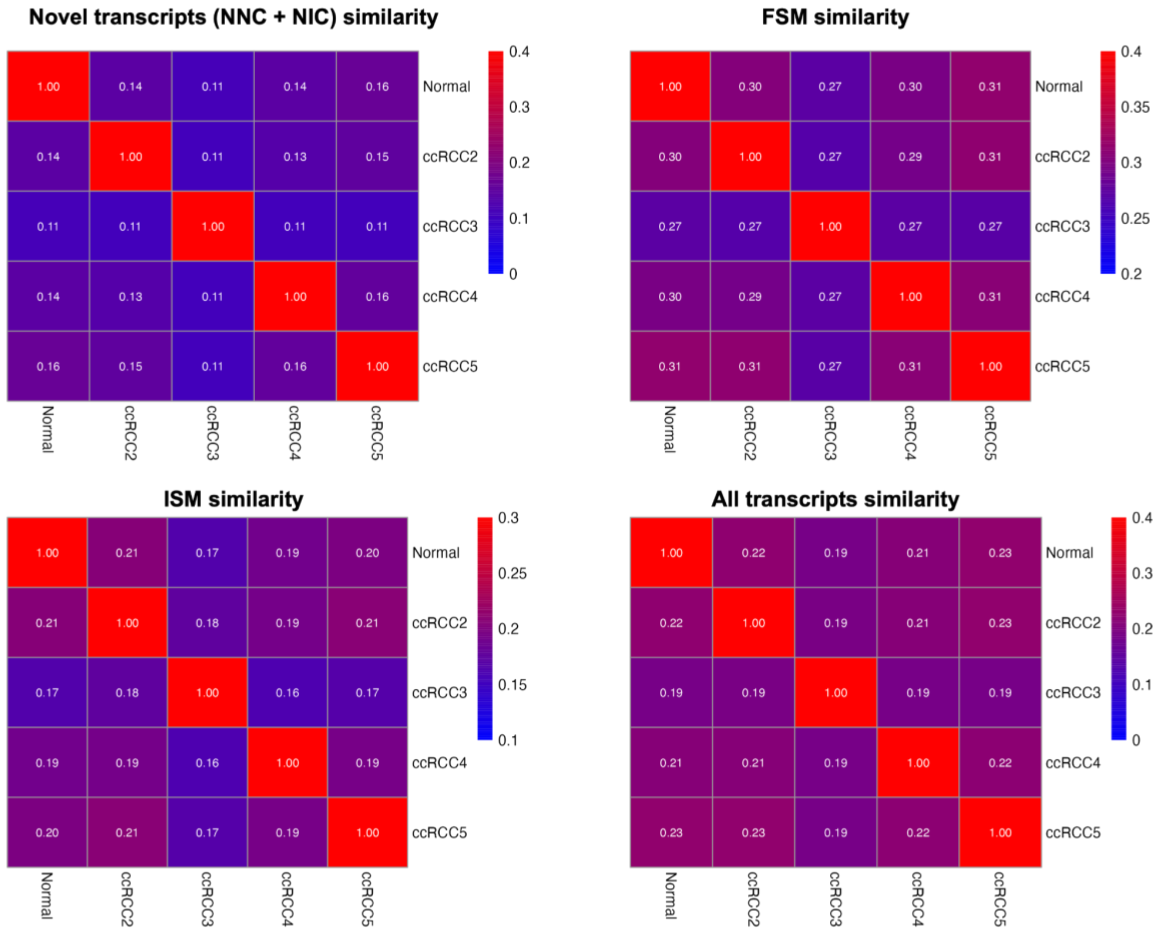


**Supplementary Figure 5: Transcript Similarity and Cell Prevalence in Samples:** Heatmaps display the Jaccard similarity index among samples for different categories of transcripts. Top left: Novel In catalog and novel not in catalog transcripts (NIC and NNC, respectively) similarity; top right: Full Splice Match (FSM) similarity; bottom left: Incomplete Splice Match (ISM) similarity; bottom right: Overall similarity including all transcripts, with the criterion that transcripts must be present in at least three cells.

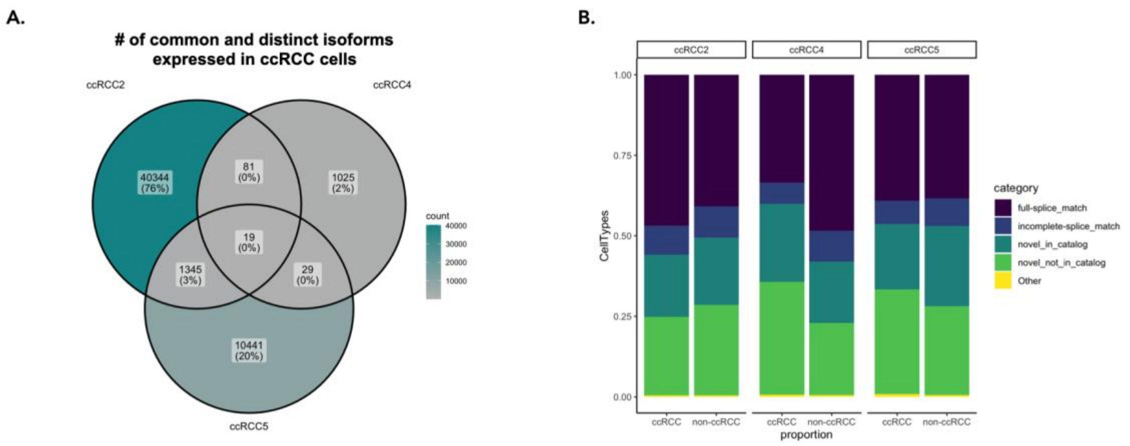


**Supplementary Figure 6: (A).** Number of overlapping isoforms explicitly in ccRCC cells. **(B).** Proportion of explicitly expressed isoforms either in ccRCC or non-ccRCC cells across ccRCC2, ccRCC4, ccRCC5.

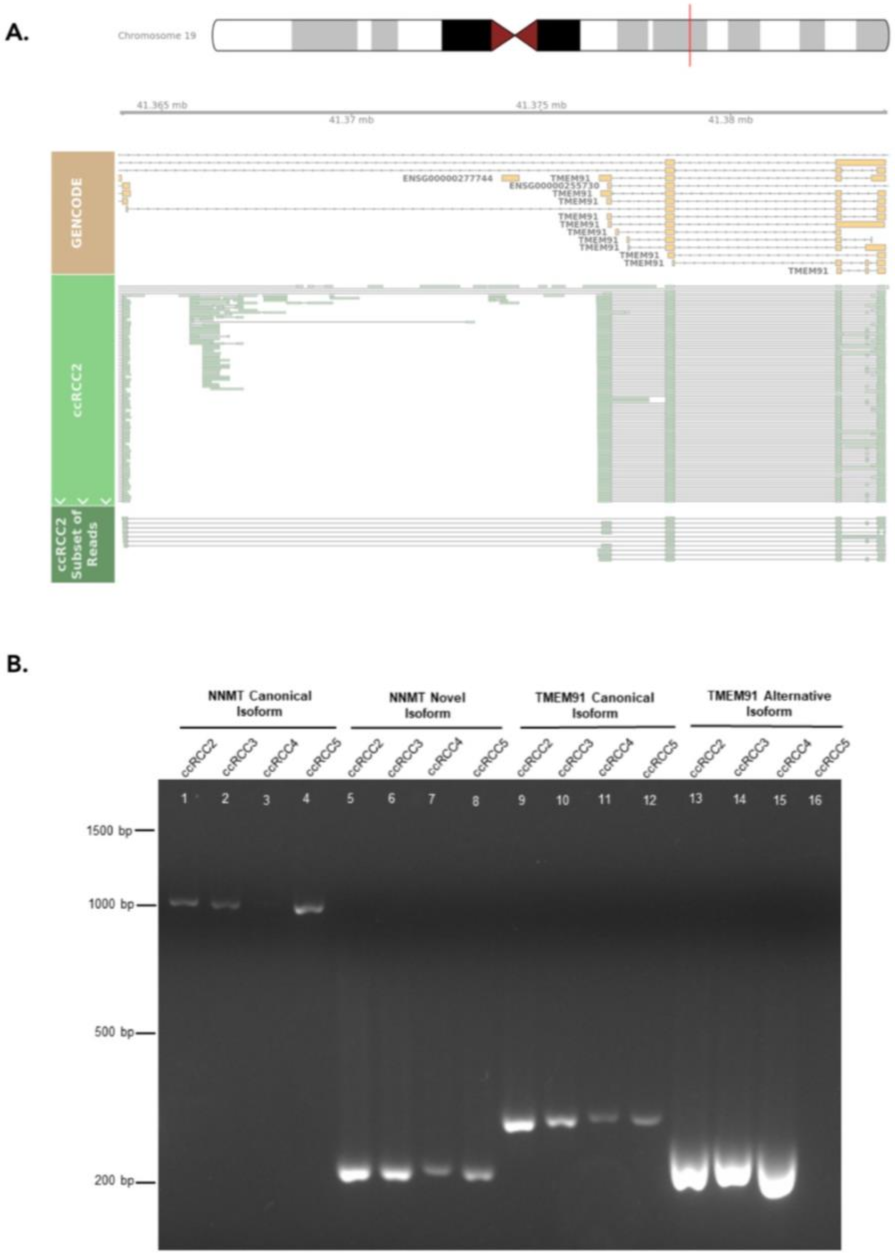


**Supplementary Figure 7: (A).** TMEM91 novel isoform reads. 520 PacBio reads were found for the transcripts. **(B). PCR Validations of Isoforms in ccRCC Tumor Samples:** Agarose gel (2%) electrophoresis image of PCR products amplified with common or isoform specific primers (described in supplementary table 1 & 2)**.** All lanes are marked with corresponding tumor sample names. Lane 1-4: NNMT Canonical Isoform amplified with NNMT_Canonical_Fp and NNMT_Common_Rp1, lane 5-6: NNMT Novel Isoform amplified with NNMT_Novel_Fp and NNMT_Common_Rp2, lane 9-12: TMEM91 Canonical Isoform amplified with TMEM91_Common_Fp and TMEM91_Common_Rp, lane 13-16: TMEM91 Alternative Isoform amplified with TMEM91_Novel_Fp and TMEM91_Common_Rp. The novel transcript included five exons without exon 2. In both PCR and Sanger sequencing, we could confirm the exon 1 exon 3 junction of transcript ENST00000413014. Note that the target novel isoform was lost after filtering, which may explain its absence in the PCR results**.**

**Supplementary Table 1: Details of Primers used in PCR Validation**

| Target | Primer specification | Primer name | Sequence |
|---|---|---|---|
| NNMT | Specific forward primer against unique sequence of canonical isoform | NNMT_Canonical_Fp | TCAAGTGCTCCCTCTGGTCT |
| | Specific forward primer against unique sequence of novel isoform | NNMT_Novel_Fp | TGGTGTCTACTTCTTGGCTTTTG |
| | Reverse primer against sequence shared between canonical and novel isoforms | NNMT_Common_Rp1 | ACCGCCTGTCTCAACTTCTC |
| | Reverse primer against sequence shared between canonical and novel isoforms | NNMT_Common_Rp2 | AGTAGGTGGGGAGGTCTGG |
| TMEM91 | Forward primer against shared sequence between canonical and novel isoforms | TMEM91_Common_Fp | AGACCCGCGTAGAGCAAAG |
| | Specific forward primer against unique sequence of novel isoform | TMEM91_Novel_Fp | GGGCTTGACTGCTTCTTTTC |
| | Reverse primer against sequence shared between canonical and novel isoforms | TMEM91_Common_Rp | CTCGGCAAAGGCTATCTCTC |

**Supplementary Table 2: Details of Anticipated and Observed Outcome of PCR Validations Using Different Primer Pairs**

| Target | | Primer Combination | Anticipated Outcome | Observed Outcome |
|---|---|---|---|---|
| NNMT | NNMT Canonical | NNMT_Canonical_Fp | Single PCR product corresponding to canonical isoform | Matched with anticipated outcome |
| | | NNMT_Common_Rp1 | | |
| | NNMT Novel | NNMT_Novel_Fp | Single PCR product corresponding to novel isoform | Matched with anticipated outcome |
| | | NNMT_Common_Rp2 | | |
| TMEM91 | TMEM91 Common | TMEM91_Common_Fp | Single PCR products corresponding to both canonical and novel isoforms | Matched with anticipated outcome |
| | | TMEM91_Common_Rp | | |
| | TMEM91 Novel | TMEM_Novel_Fp | Single PCR product corresponding to novel isoform | Single PCR product but with shorter length |
| | | TMEM_Common_Rp | | |

**Supplementary Table 3: Comparison of cMDT calculations and Acorde results.** The table shows the overlap of upregulated isoforms in non-ccRCC cells of ccRCC2 found by Acorde and MDTs in non-ccRCC cells of ccRCC2. Isoforms found by two methods are highlighted in bold.

| cMDT Tama ID | Gene Name | Cell Number | MDT in Normal | Cohort | Acorde Result |
|---|---|---|---|---|---|
| G14020.26 | ENSG00000237550 | 4 | **G14020.22** | ccRCC2 | **G14020.22** |
| G16060.56 | CYBA | 1 | **G16060.25** | ccRCC2 | **G16060.25** |
| G42843.68 | JPX | 4 | G42843.13, G42843.130, **G42843.150**, G42843.23, G42843.25 | ccRCC2 | **G42843.150** |
| G42843.1 | JPX | 3 | G42843.13, G42843.130, **G42843.150**, G42843.23, G42843.25 | ccRCC2 | **G42843.150** |
| G42843.75 | JPX | 3 | G42843.13, G42843.130, **G42843.150**, G42843.23, G42843.25 | ccRCC2 | **G42843.150** |
| G42843.107 | JPX | 1 | G42843.13, G42843.130, **G42843.150**, G42843.23, G42843.25 | ccRCC2 | **G42843.150** |
| G42843.127 | JPX | 1 | G42843.13, G42843.130, **G42843.150**, G42843.23, G42843.25 | ccRCC2 | **G42843.150** |
| G42843.145 | JPX | 1 | G42843.13, G42843.130, **G42843.150**, G42843.23, G42843.25 | ccRCC2 | **G42843.150** |
| G42843.148 | JPX | 1 | G42843.13, G42843.130, **G42843.150**, G42843.23, G42843.25 | ccRCC2 | **G42843.150** |
| G42843.159 | JPX | 1 | G42843.13, G42843.130, **G42843.150**, G42843.23, G42843.25 | ccRCC2 | **G42843.150** |

| | | | | | |
|---|---|---|---|---|---|
| G42843.212 | JPX | 1 | G42843.13, G42843.130, **G42843.150**, G42843.23, G42843.25 | ccRCC2 | **G42843.150** |
| G42843.35 | JPX | 1 | G42843.13, G42843.130, **G42843.150**, G42843.23, G42843.25 | ccRCC2 | **G42843.150** |
| G42843.44 | JPX | 1 | G42843.13, G42843.130, **G42843.150**, G42843.23, G42843.25 | ccRCC2 | **G42843.150** |
| G42843.56 | JPX | 1 | G42843.13, G42843.130, **G42843.150**, G42843.23, G42843.25 | ccRCC2 | **G42843.150** |
| G42843.64 | JPX | 1 | G42843.13, G42843.130, **G42843.150**, G42843.23, G42843.25 | ccRCC2 | **G42843.150** |
| G42843.81 | JPX | 1 | G42843.13, G42843.130, **G42843.150**, G42843.23, G42843.25 | ccRCC2 | **G42843.150** |
